# A High-Throughput Platform for Rapid Adaptation of DNA Aptamers to SARS-CoV-2 Evolution

**DOI:** 10.64898/2026.04.21.719937

**Authors:** Yujie He, Zhenglin Yang, Yu-An Kuo, Yuting Wu, Diego Fonseca-Albert, Kyle K. Le, Jeffrey Guo, Yanxing Wang, Anh-Thu Nguyen, Yuan-I Chen, Sohyun Kim, Wei-Ru Chen, Saeed Seifi, Soonwoo Hong, Trung Duc Nguyen, Yinong Chen, Pengyu Ren, Yi Lu, Hsin-Chih Yeh

## Abstract

Rapid pathogen evolution threatens public health by eroding the effectiveness of vaccines, therapeutics, and diagnostic tools. Although spike protein targeting monoclonal antibodies (mAbs) were developed within 10-12 months of the initial outbreak to serve as key theranostic agents, their redesign has struggled to keep pace with viral evolution, rendering many neutralizing antibodies ineffective. Here we demonstrate a novel platform that combines a random-rational hybrid library diversification with high-throughput *MiSeq* screening to rapidly reprogram aptamers against emerging SARS-CoV-2 spike variants. Interactions between 3 different spike proteins and 11,806 unique aptamer variant designs were profiled within a few days. Starting from a 40-nt aptamer originally selected against wild-type (WT) spike protein, our screen identified a Delta-binding mutant with a 4-fold affinity improvement and an Omicron-binding mutant that converted undetectable binding into nanomolar affinity. We also identified a WT-selective mutant with substantially reduced affinity for Delta, as well as previously unrecognized bases that critically contribute to spike recognition. Integrating high-throughput binding data with molecular dynamics simulations further revealed sequence-dependent structural features underlying variant-specific aptamer-spike interactions. Finally, we developed a sensor based on the identified WT-selective aptamer mutant, enabling highly specific detection of the WT spike protein with robust performance. Together, this work establishes a rapid and adaptable aptamer engineering platform for rapid adaptation of aptamers to evolving pathogens in future pandemics.

## INTRODUCTION

By December 2025, the cumulative number of confirmed COVID-19 cases and deaths worldwide had reached 779 million and 7.1 million, respectively^1^. A major challenge in halting this pandemic arises from the rapid evolution of SARS-CoV-2 virus. Since the outbreak in 2019, the World Health Organization (WHO) has identified five variants of concerns (VOCs), including Alpha, Beta, Gamma, Delta and Omicron^2–6^, These variants exhibit markedly higher transmissibility than the original strain^7,8^, leading to extended quarantine, increased hospitalization and vaccine breakthrough infections^9–15^. To combat the COVID-19 pandemic, at least 17 monoclonal antibodies (mAbs) for both patient treatment and SARS-CoV-2 detection were successfully developed and granted Emergency Use Authorization (EUA) by the U.S. FDA in late 2020^16,17^. However, these authorizations were later revoked^18,19^ as continuous viral evolution rendered these antibodies ineffective^20,21^. Developing new mAbs at pandemic speed and subsequently optimizing them to keep pace with viral mutations^18,20^ remain a formidable challenge. Therefore, a modular, highly adaptable platform that can be rapidly reprogrammed and deployed for both the diagnosis and therapeutic applications of emerging viral variants is urgently needed to support rapid responses for future pandemics^22^.

Highly adaptable functional nucleic acid technologies can fulfill this need. Analogous to mAbs, DNA aptamers identified through SELEX (<u>S</u>ystematic <u>E</u>volution of <u>L</u>igands by <u>EX</u>ponential enrichment) process^23–25^ can bind spike proteins with high affinities (*k*_*d*_ values down to the sub nM range^26–28^). For instance, a 40-nt DNA aptamer Apt2 developed by Zu and co-workers^27^ exhibits a *k*_*d*_ of 2 nM when binding spike protein derived from the original wild-type (WT) strain and costs only ∼$0.72 per nanomole to synthesize commercially. In the past few years, dozens of DNA aptamers have been developed to bind different epitopes of the SARS-CoV-2 virus, such as various domains, especially the receptor-binding domain (RBD), of the spike protein^26,28–31^ and nucleocapsid proteins^32^. Although some can bind multiple viral strains^33–35^, most of them faced limited cross-variant robustness. For instance, we found that Apt2 exhibits negligible binding to the spike protein of one of the Omicron subvariants, Omicron_XBB_ (**Fig. 1C**), possibly because Apt2 was originally selected against the WT strain, whereas the subvariant harbors 39 amino-acid mutations in its spike protein (21 of them are within the RBD). Although aptamers, compared to mAbs, are inherently more amenable to rapid reengineering to accommodate viral mutations, the conventional de novo approach to develop DNA aptamers (i.e., SELEX^23^) relies on time-consuming iterative enrichment and counterselection steps to isolate strong and highly specific binders from a highly diverse library (∼10^15^ sequences), limiting its ability to generate new aptamers quickly enough to keep pace with viral evolution during a pandemic.

**Fig. 1.**
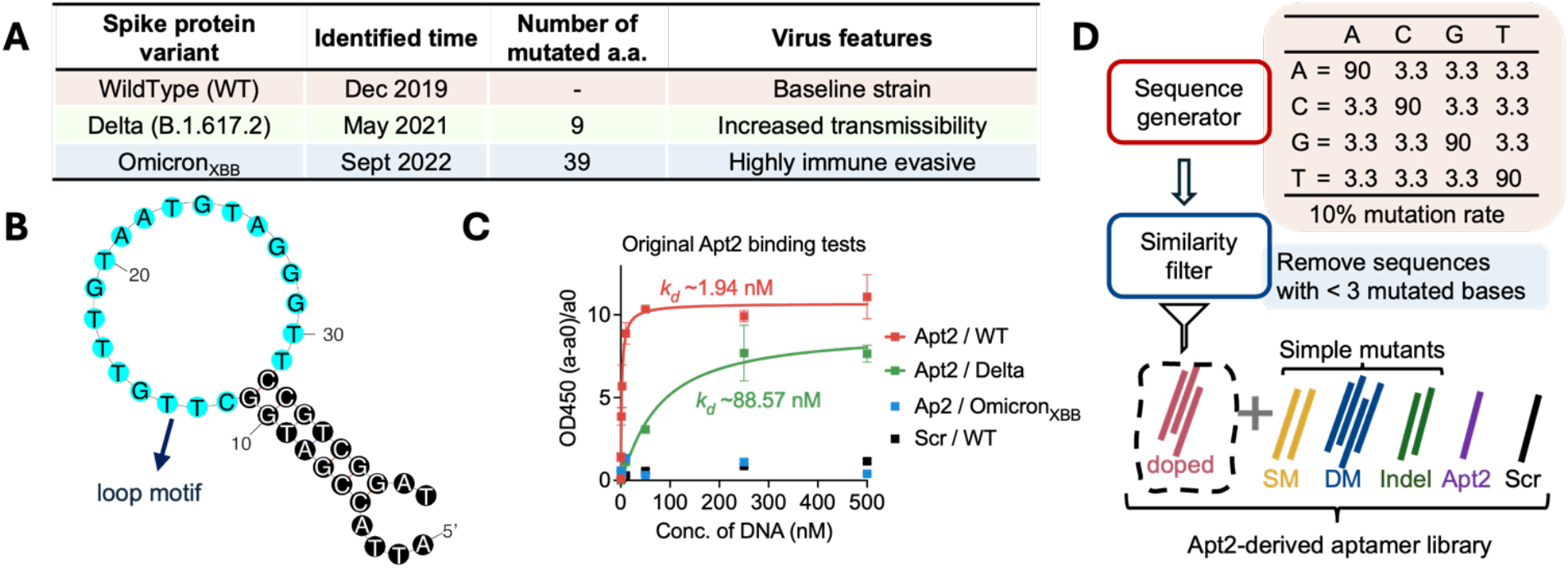
Apt2 chosen as the seeding strand for SARS-CoV-2 spike protein aptamer diversification and screening. (**A**) The selected SARS-CoV-2 spike protein (trimer) variants in the experiment. Both Delta and Omicron_XBB_ drove major global waves during the COVID-19 pandemic through their mutations, with the more recent one Omicron_XBB_ being distinctly different than the original strain. a.a.: amino acid(s). (**B**) The secondary structure of Apt2 as predicted by mFold, with the 20-nt loop motif region identified by the previous SELEX, highlighted in light blue. (**C**) Characterization of the seeding aptamers and the negative control sequence with ELONA. *k*_*d*_(Apt2/WT) ∼2±0.4 nM; *k*_*d*_(Apt2/Delta) ∼89±10 nM. (**D**) Aptamer library design for chip screening. SM: single mutants; Indel: insertion and deletion mutants; DM: double mutants; Scr: the scrambled sequence.

To overcome this limitation, we developed a pipeline based on a repurposed Illumina next-generation sequencing (NGS) platform^36–50^ that enables massively parallel, quantitative evaluation of aptamer–protein interactions across 10^4^-10^7^ (depending on the flow cell type) closely related sequences within a single run^51–55^. Repurposed NGS flow cells have provided a powerful platform to directly study interactions between target analytes and libraries of DNA^40,53^, RNA^45,46^, modified bases^56^, and peptide^42,47^. Previously, we employed a repurposed NGS method known as CHAMP^53^ (<u>C</u>hip-<u>H</u>ybridized <u>A</u>ssociation-<u>M</u>apping <u>P</u>latform, which eliminates the need to modify Illumina’s sequencing protocol) to screen fluorescent nanomaterial^57^, optimize and diversify the photophysical properties of fluorogenic DNA^55^ and RNA^58^ aptamers, while related studies^56,59–61^ have demonstrated the broader utility of NGS-based platforms for aptamer characterization and optimization. While previous work established a high-throughput screening framework for nanomaterial characterization, its ability to rapidly adapt to the emerging viral variants remains limited.

To address this, we developed a random-rational hybrid library design (**Fig. 1D**) integrated with CHAMP-based screening. This approach is based on the premise that, for certain virus-binding aptamers, adaptation to new variants does not require complete redesign, but can be achieved through targeted diversification around a functional seeding sequence. By preserving controlled diversity in this local sequence space, the platform enables rapid and directed aptamer reengineering without the need for de novo selection. Using this strategy, we screened 11,806 variants directly on a repurposed NGS flow cell. Apt2 served as an ideal model due to its well-characterized WT RBD binding and early development during the pandemic. By simultaneously measuring binding affinities across all variants without enrichment or loss of sequence diversity, this platform enables systematic exploration of sequence-function relationships and accelerates adaptation to new SARS-CoV-2 variants from months to days. The reusability of the flow cell further supports iterative testing against multiple viral variants on the same platform, facilitating rapid identification of both affinity-enhanced and strain-selective binders. To the best of our knowledge, this study is the “first to demonstrate repurposed NGS screening as a rapid adaptation strategy for reengineering aptamer recognition against evolving pathogen variants.

In this study, we show that this high-throughput screening strategy enables rapid reengineering of aptamers to adapt to emerging SARS-CoV-2 variants from a single WT binder. Specifically, targeted diversification yielded variants with enhanced binding to Delta, restored affinity toward Omicron_XBB_ from undetectable levels to the nM range and enabled the generation of WT-selective binders with minimal cross-reactivity. The high-throughput profiling also revealed that beyond the loop motif, the upper stem region plays a critical role in spike protein recognition. Further molecular dynamics (MD) simulations offered structural rationales for the improved binding characteristics of the optimized aptamer mutants. Finally, a WT-selective aptamer was translated into a strand-displacement fluorescent sensor, achieving highly specific detection with robust performance in complex biological matrices. Together, these results demonstrate a scalable strategy for rapidly adapting molecular recognition elements to evolving pathogens, providing a practical framework for next-generation theranostic development.

## RESULTS

### Designing an Apt2 Variant Pool for Aptamer Diversification

We selected Apt2, a 40-nt DNA aptamer developed by the Zu group^27^ against the SARS-CoV-2 WT spike protein (hereafter referred to as WT in short), as the seeding aptamer to explore neighboring sequence variants capable of binding the Delta (B.1.617.2) and Omicron_XBB_ spike proteins. Apt2 was chosen because of its well-characterized WT RBD binding and its early development during the pandemic. Delta and Omicron_XBB_ were selected as model viral variants because both raised major global concern and complicated pandemic control through increased transmissibility and reduced immune effectiveness^5,13–15,20^. They also represent distinct stages and magnitudes of SARS-CoV-2 evolution, with Delta and Omicron_XBB_ carrying 9 and 39 spike amino-acid mutations, respectively, and emerging approximately 1 and 3 years after the original strain (**Fig. 1A**). Together, these features make Apt2 and these two variants suitable models for evaluating whether local sequence diversification can rapidly adapt an existing aptamer to evolving viral targets.

The Apt2 folds into a hairpin structure, as predicted by mFold^62^, with the 20-nt loop corresponding to the binding motif identified through the SELEX process (**Fig. 1B**). We first tested the binding affinity of the original Apt2 against the aforementioned three spike proteins using an enzyme-linked oligonucleotide assay^63^ (ELONA, **fig. S1A**). The results (**Fig. 1C**) were consistent with the structural and sequence divergence of these spike proteins, showing that Apt2 binds strongly to the WT (*k*_*d*_ ∼2±0.4 nM), moderately to the Delta spike protein (*k*_*d*_ ∼89±10 nM) and becomes undetectable with the Omicron_XBB_ spike protein (no stable *k*_*d*_ determined). No binding affinity to any of the three spike proteins was observed from the scramble sequence (Scr). A mutant pool was designed using a random-rational hybrid algorithm to ensure a controlled diversity in this local sequence space, which included four subgroups: single mutations (120 sequences, 1.08%), double mutations (6,508 sequences, 58.7%), indels (single insertions or deletions, 152 sequences, 1.37%), and doped (i.e., partially randomized^64,65^) mutants (4,500 sequences, 37.6%, generated with a mutation rate of 10% within the 20-nt loop motif and filtered to remove duplicates with the other subgroups) (**Fig. 1D**). Based on the calculation (**Methods** and **fig. S2**), most doped mutants contain either triple (59%) or quadruple (28%) mutations, representing a well-controlled degree of diversity for optimizing aptamers for emerging variants.

### High-throughput Screening of Apt2-derived Pool on an Illumina NGS Flow Cell

To screen the pool in a high-throughput manner, we synthesized the library, added sequencing adapters and loaded it onto an Illumina *MiSeq* flow cell (**Fig. 2A**). Before screening, we first confirmed that the sequencing adapters and primers did not disrupt aptamer binding (**figs. S3-S4**). After sequencing, the strands were denatured, and the flow cell was thoroughly rinsed to remove any excess dyes. The fiducial marker PhiX was then labeled with an Atto488-tagged PhiX probe and imaged (**Fig. 2B**) for mapping purpose^51,55^. The His-tagged spike protein was injected to the flow cell, captured by the displayed binders, and secondarily labeled via an Alexa647-conjugated anti-His-tag antibody. The fiducial markers and bound proteins were imaged under fluorescence microscopy using two different filter cubes). The spike protein binding capacity of each sequence was estimated by the mean fluorescence intensity among its duplicates on the chip (**Methods**). By digesting the proteins and denaturing the DNAs, the same flow cell could be reset to its initial state for sequential screening of binders against another protein target (**Fig. 2C**). This approach allows the binding characterization of the entire pool against multiple SARS-CoV-2 spike protein variants on a single chip, thereby reducing experimental cost and material usage.

**Fig. 2.**
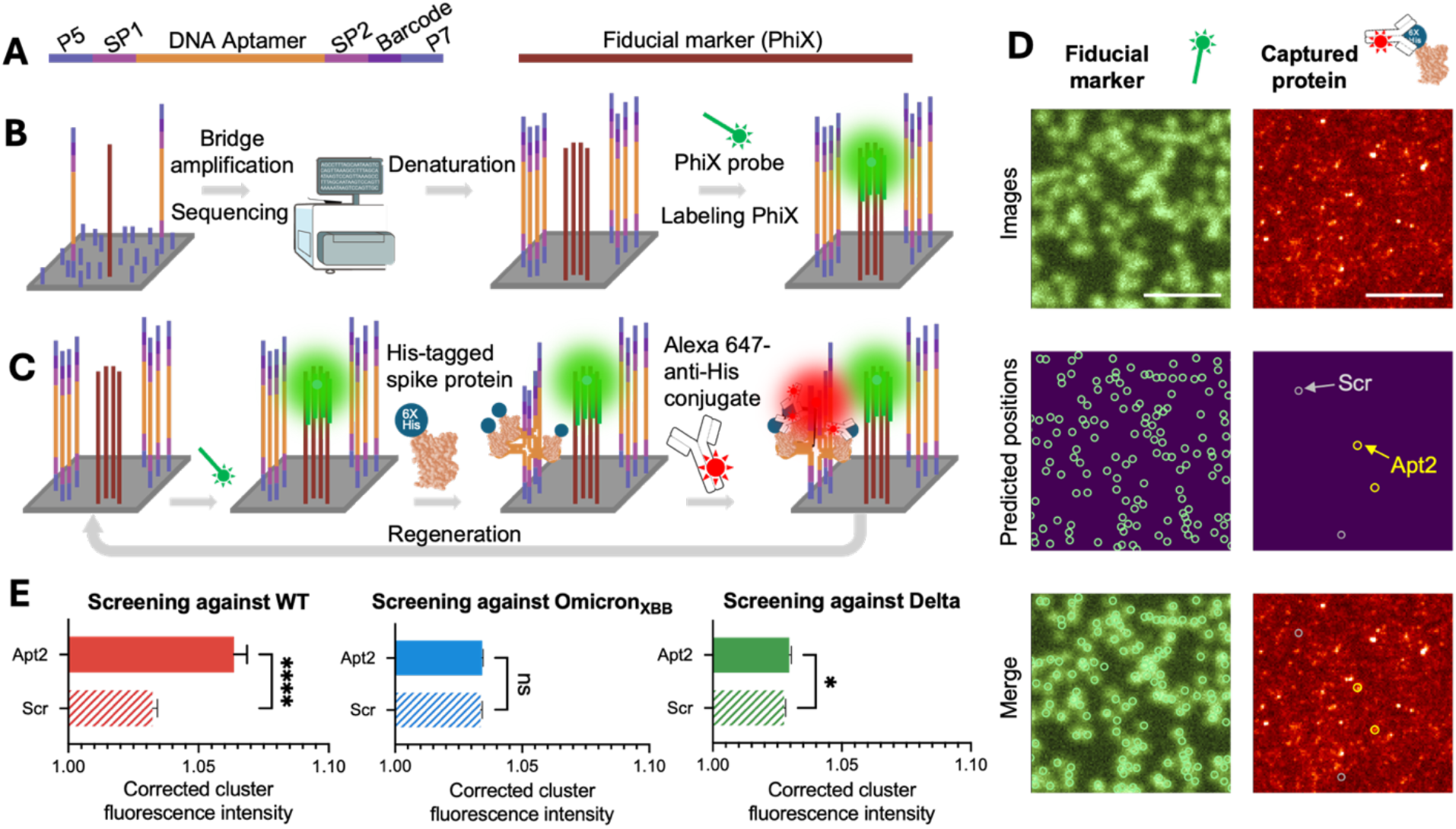
Overview of SARS-CoV-2 spike protein variants specific aptamer identification on Illumina NGS chips. (**A**) The DNA pool described in **Fig. 1** are sandwiched in between Illumina sequencing adapters and synthesized on the flow cell surface. (**B**) NGS library was sequenced on a *MiSeq* chip through standard PE75 protocol and was denatured to expose ssDNA on the chip surface. Control sequences (i.e., PhiX) were labeled with an Atto488-tagged probe (fiducial marker). (**C**) To image SARS-CoV-2 spike protein binding on the chip, we first incubated the flow cell with 100 nM His-tagged spike protein trimer, which was then stained by the Alexa647 His-tag antibody conjugate. After imaging, the chip was regenerated by 0.1 N NaOH treatment, with another spike protein variant added in the following round. (**D**) A representative field of view (FOV) of PhiX labeled with Atto488-tagged probes (left, top) for mapping purpose, and WT spike protein labeled with Alexa647-His-tag antibody conjugate (right, top). The polonies representing the PhiX sequences, the seeding aptamer Apt2 and the negative control sequence (Scr) are highlighted in green (left, middle), yellow and light gray (right, middle) circles, respectively, which were identified after the CHAMP alignment. The merged figures (bottom) show the successful mapping of sequence positions and the chip images. Scale bars are 10 µm. (**E**) Corrected fluorescence intensities of the polonies correlating the seeding aptamers and the negative control sequence (Scr) on the chip when co-incubated with WT, Omicron_XBB_, and Delta spike proteins, respectively. Error bars reflect the SEM (standard error of the mean) of each group. Number of duplicates of each trial can be found in **Supplementary Table S3.** *, p < 0.05; **, p < 0.01; ***, p <0.001; ns, p > 0.05.

As a proof-of-concept, we performed the first screen against WT. Fiducial markers’ patterns were captured under the FITC channel, and those of captured spike proteins were collected under the Cy5 channel (**Fig. 2D, upper**). The positions of the bright FITC puncta matched the coordinates recorded in the FASTQ file (**Fig. 2D, left**), confirming successful image alignment. Within the field of view (FOV), positions corresponding to Apt2 co-localized with several bright Alexa647 puncta, whereas those corresponding to Scr remained dark (**Fig. 2D, right**). These results demonstrated that binding was visualized as planned. Next, we regenerated the flow cell and performed another round of screening against Omicron_XBB_. We compared the acquired fluorescence intensities with the original Apt2 and Scr signals. The results showed that Apt2 exhibited significantly higher fluorescence intensity than Scr when introducing WT (**Fig. 2E, left**), while the brightness of them was similar when the protein target switched to the Omicron_XBB_ (**Fig. 2E, middle**), which agreed with our ELONA results that Apt2 is a strong binder to the WT while a non-binder to the Omicron_XBB_ (**Fig. 1C**). A third round of screening against Delta was then performed following the chip regeneration (**Fig. 2E, right**).

### Enhancing the Affinity for Delta by Modifying the Aptamer Stem

Here we sought to identify mutants with improved binding to the Delta strain, as a proxy for adaptation to an earlier stage of SARS-CoV-2 evolution. We quantified the percentage change in fluorescence signal for each sequence relative to original Apt2 (hereafter referred to as percentage change) across three independent screening trials, each performed against a different strain. Analysis of the combined WT and Delta screening data showed that most sequences showed significantly lower binding to WT compared to Apt2, while approximately half pool exhibited enhanced binding to Delta (**fig. S5**). Nevertheless, because the fluorescence signals of Apt2 and Scr were less clearly separated for Delta (**Fig. 2E**), the corresponding calculated percentage changes could be less reliable. This limitation was further revealed by streptavidin-mediated surface binding assays (**fig. S1B**), where several high-fluorescence candidates failed to show consistently enhanced binding in ELONA (e.g., T3C-A39C; **Fig. 3C** and **fig. S6**).

**Fig. 3.**
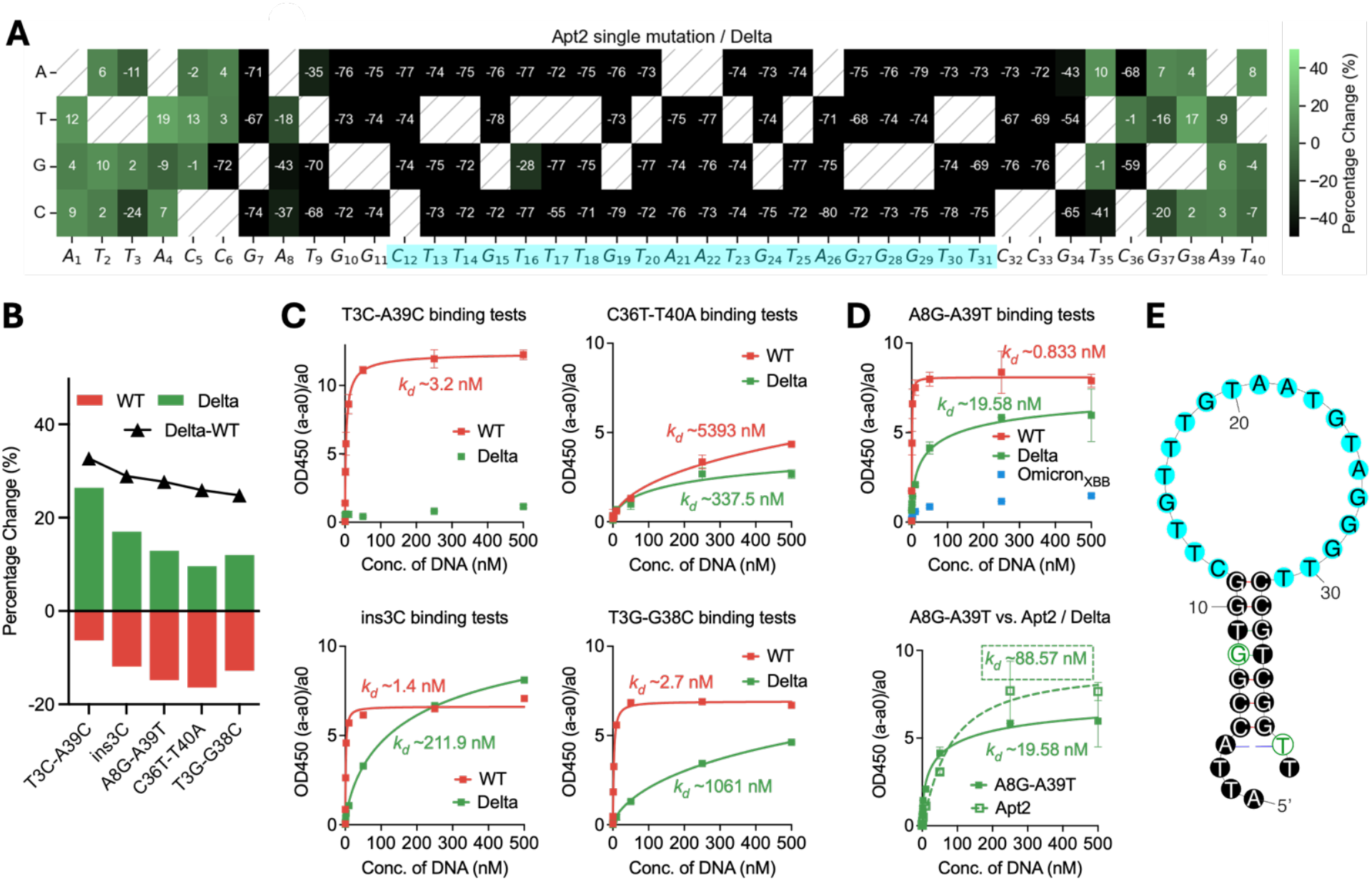
Identifying Apt2-A8G-A39T as an improved Delta binder. (**A**) Apt2 single-mutation/Delta heatmap. The color of each cell represents the percentage change compared to the original Apt2/Delta with the color bar showing on the right. Cells representing the unmutated sequences are shown as diagonally hatched. The heatmap implied the immutability of the loop motif (highlighted in cyan). (**B**) Top five stem-altered candidates ranked by the Delta-WT percentage-change difference. Percentage changes for WT and Delta are shown as red and green bars, respectively. The black line indicates the Delta-WT percentage-change difference. (**C**)-(**D**) ELONA revealed that among all 5 candidates, A8G-A39T can bind slightly stronger than the seeding aptamer Apt2 (*k*_*d*_ ∼20±4 nM vs. *k*_*d*_ ∼89±10 nM, right). Further investigation showed that A8G-A39T has a similar binding pattern to the original Apt2, which is a strong WT binder (*k*_*d*_ ∼0.8±0.2 nM), medium Delta binder, and does not bind Omicron_XBB_. (**E**) A8G and A39T substitutions (highlighted by the green circles) were highlighted in the mFold-predicted secondary structure of Apt2. The motif was highlighted in cyan.

To overcome the challenge, we changed the strategy by examining binding patterns at the sequence level using the screening data. The effects of individual sequences on a given viral strain were summarized as percentage changes and visualized across a series of heatmaps. Single mutants (**Fig. 3A**) and indels (**fig. S7**) binding data against Delta revealed that the 20-nt loop motif (residues C12 to T31) was also conserved for Delta binding, since any point mutation or insertion within this region disrupted interaction. Double-mutation heatmap on the stem further showed that the bases close to the loop are also intolerant with mutations (**fig. S8**). Based on this observation, we restricted subsequent analysis to mutants with alterations confined to the stem region to preserve Delta binding. These candidates were ranked according to the difference in percentage change between WT and Delta, with the goal of minimizing cross-reactivity. Top five sequences were selected for further evaluation (**Fig. 3B**). The binding profiles were validated outside the chip using ELONA (**Fig. 3C**). Most candidates exhibited binding profiles similar to that of the original Apt2. Among them, A8G-A39T showed a modest improvement in binding affinity for Delta relative to Apt2 (*k*_*d*_ ∼20±4 nM vs. 89±10 nM). Notably, this variant retained very strong binding to WT (*k*_*d*_ ∼0.8±0.2 nM) and negligible affinity for Omicron_XBB_ (**Fig. 3D**).

### Identification of Omicron_XBB_ Binders

To assess whether our platform can identify strands targeting more recent variants, we screened the designed pool for Omicron_XBB_ binders. Most fluorescence signals were comparable to those of Apt2 and Scr (**Fig. 4A**), indicating that the majority exhibit weak or no binding to Omicron_XBB_. Notably, a small subset displayed a pronounced increase in fluorescence intensity and was selected for further investigation. To improve binding specificity and reduce background from multi-target screens, we narrowed the selection to five candidates with negligible signals in both the WT and Delta screens (**Fig. 4B**, highlighted in blue). Among these, two were double mutants with the mutated bases positioning within the 20-nt loop motif, while three originated from the doped mutation subgroup (**Fig. 4B**, zoom-in), suggesting that unlike WT and Delta, the loop motif is insufficient to support Omicron_XBB_ binding (**figs. S11-12**).

**Fig. 4.**
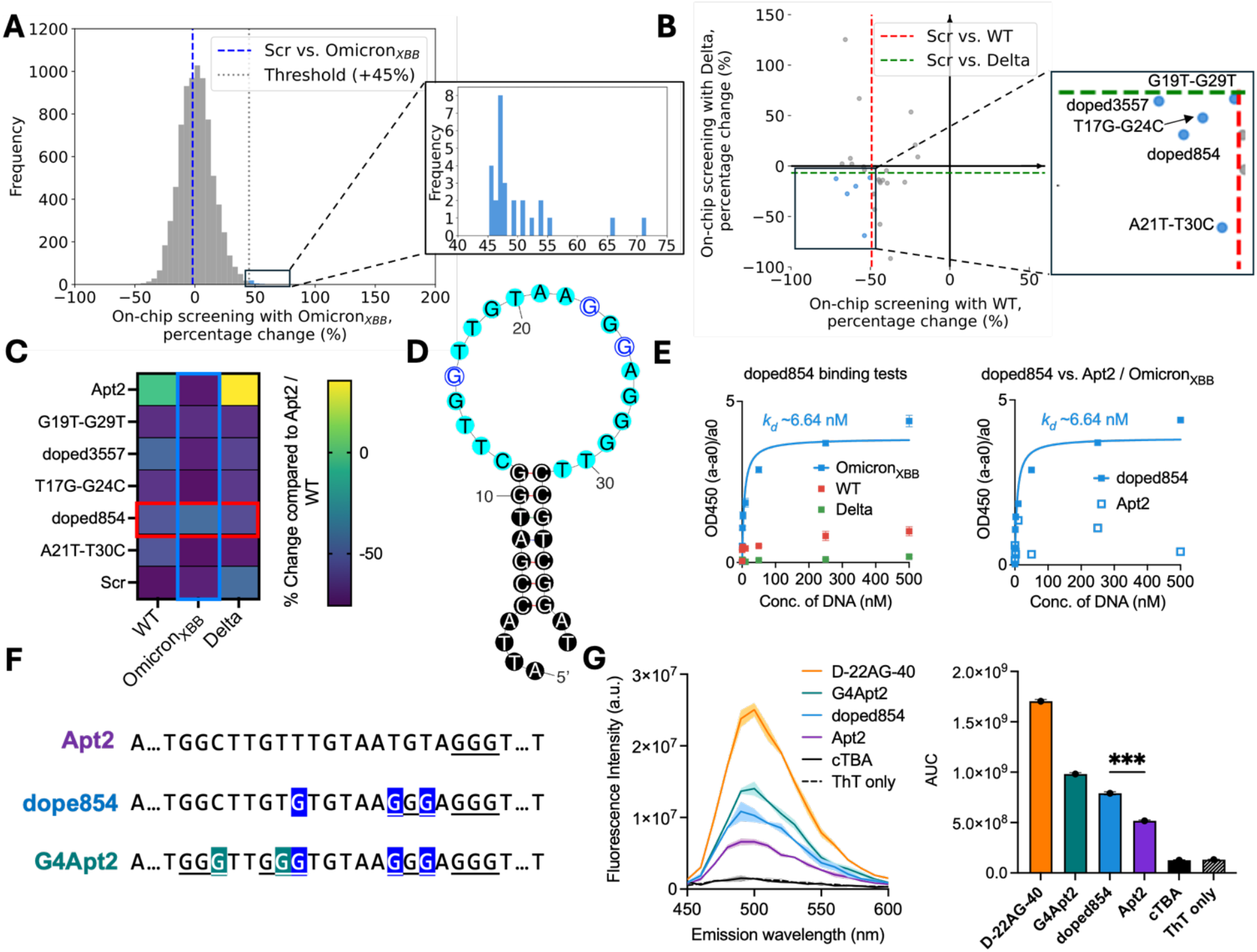
Identifying Apt2-doped854 as an OmicronXBB-selective binder. (**A**) Sequences with at least +45% higher intensity than that of Apt2/Omicron_XBB_ were selected out from the whole pool (bars in blue). **(B)** These selected candidates’ WT and Delta binding data were retrieved and summarized in a scatterplot, with those with neglect binding affinity for WT (to the left of the red dotted line) and Delta (below the green dotted line) highlighted as blue puncta. (**C**) Streptavidin-mediated surface binding assay was used to test the bindings of Omicron_XBB_ candidates selected from (**A**)-(**B**). Promising candidate doped854 was highlighted in a red box. Binding affinity for Omicron_XBB_ comparison among the candidates are highlighted in a blue box. The color scale is shown on the right. (**D**) T17G, T23G and T25G substitutions (highlighted by the blue circles) were indicated in the mFold-predicted secondary structure of Apt2. The loop motif was highlighted in cyan. (**E**) ELONA revealed that Apt2-doped854 can bind Omicron_XBB_, developing a new binding feature compared to the original Apt2 (*k*_*d*_ ∼7±0.9 nM vs. undetectable). (**F**) Sequences of Apt2, doped854 and G4Apt2. Residues T2 to A8, T31 to A39 were omitted. Mutated bases in doped854 were highlighted in blue and further altered bases in G4Apt2 were highlighted in teal. G-tracts (G3s) were underscored. (**G**) Fluorescence activation of thioflavin T by DNA strands. D-22AG-40 is a G4 dimer known for strongly enhancing ThT’s fluorescence. cTBA is the reverse complementary sequence of thrombin’s aptamer TBA, which has no guanine in the sequence. Fluorescence was measured using 1 µM DNA and 1 µM thioflavin T. Shadings represent the standard deviations. Area under curve (AUC) was used to generate the overall intensities shown in the bar chart. Two-tailed un-paired *t* test was performed to compare the fluorescence intensities between doped854 and Apt2. P-value = 0.0001.

A streptavidin-mediated surface binding assay was used to evaluate the binding profiles of the five selected sequences (**Fig. 4C**). In this assay, aptamer candidates were immobilized on strep-tavidin-coated microwell plates via a biotin moiety. His-tagged spike proteins and an anti-His-fluorophore conjugate were sequentially introduced, with washing steps between incubations (**Methods**; **fig. S1B**). Fluorescence intensities were measured using a plate reader and converted to percentage changes relative to the Apt2/WT positive baseline. Four of the five candidates suppressed binding to WT and Delta but fail to exhibit improved binding to Omicron_XBB_. In contrast, doped854, a triple mutant (T17G-T23G-T25G; **Fig. 4D**), was able to exhibit binding to Omicron_XBB_. This result was further supported by ELONA (**Fig. 4E**), which confirmed that doped854 binds Omicron_XBB_ with much higher affinity than the original Apt2 (*k*_*d*_ ∼7±0.9 nM vs. undetectable).

To investigate structural features underlying doped854’s unique binding behavior, we examined its ability to activate thioflavin T (ThT), a fluorogenic dye sensitive to G-quadruplex (G4) structures^66–70^. ThT fluorescence was measured in the presence of Apt2, doped854, a rationally designed G4-forming mutant (G4Apt2: C12G-T16G-T17G-T23G-T25G; **Fig. 4F**), and control sequences known to strongly activate (D-22AG-40 ^67,70^) or not respond to ThT (cTBA ^71,72^) at the same quantity. G4Apt2 was designed based on the canonical G-quadruplex motif (G_≥3_N_x_G_≥3_N_x_G_≥3_N_x_G_≥3_)^73^. All three aptamer mutants enhanced ThT fluorescence, with doped854 producing a 53% stronger signal than Apt2 in the screening buffer and G4Apt2 yielding an even larger increase (∼90%). As expected, all signals were lower than that of D-22AG-40, a known G4 dimer (**Fig. 4G**). These results suggested that Apt2 does not form, or only weakly stabilizes, a G-quadruplex structure. Conversely, doped854 can form, or stabilize such a topology^74^, which contributes to its ability to bind Omicron_XBB_.

### Enhancement of the Binding Specificity for the Wild-Type Strain by Mutation

Although Apt2 was originally selected against WT, it also binds the Delta strain (**Fig. 2B**). Beyond improving binding affinity, we attempted to optimize specificity toward WT, with the goal of identifying aptamers that preferentially recognize WT while minimizing cross-reactivity with Delta. By isolating WT-specific binders, we aimed to improve the precision of diagnostic tools. Highly specific detection of WT provides unambiguous strain assignment, which is essential for generating strain-resolved data to support studies of viral evolutionary pathways^75^.

To identify potential sequence features associated with WT specificity, we compared the single-mutation heatmaps of Apt2 for WT (**fig. S9A**) and Delta (**Fig. 3A**), and found out that the G11 position, located at the junction between the loop and stem, tolerates a G to A substitution when binding WT, but this same mutation markedly disrupts binding to Delta (**Fig. 5A**). Consistent with this observation, the G11A single mutant showed minimal changes in binding Omicron_XBB_ (**Fig. 5B**; +18%, +7%, and -75% for WT, Omicron_XBB_, and Delta, respectively, compared to the original Apt2). WT double-mutation heatmap displayed a reddish horizontal stripe and a corresponding vertical stripe within the predominantly negative (dark) region (**fig. S10**), corresponding to the mutants where G11A is paired with a second mutated base either upstream or downstream of position 11. Notably, these cells with less negative values did not extend broadly across the heatmap, suggesting that retention of WT binding requires the second mutated base to be positioned away from the loop motif. Together, we hypothesized that Apt2 double mutants carrying G11A, combined with a second substitution outside the loop motif, might serve as selective WT binders. *MiSeq*-screening results for these double mutants are summarized in a series of bar plots (**Fig. 4C**). As expected, Delta binding was disrupted for all such double mutants, while WT binding varied depending on the type and position of the second mutated base.

**Fig. 5.**
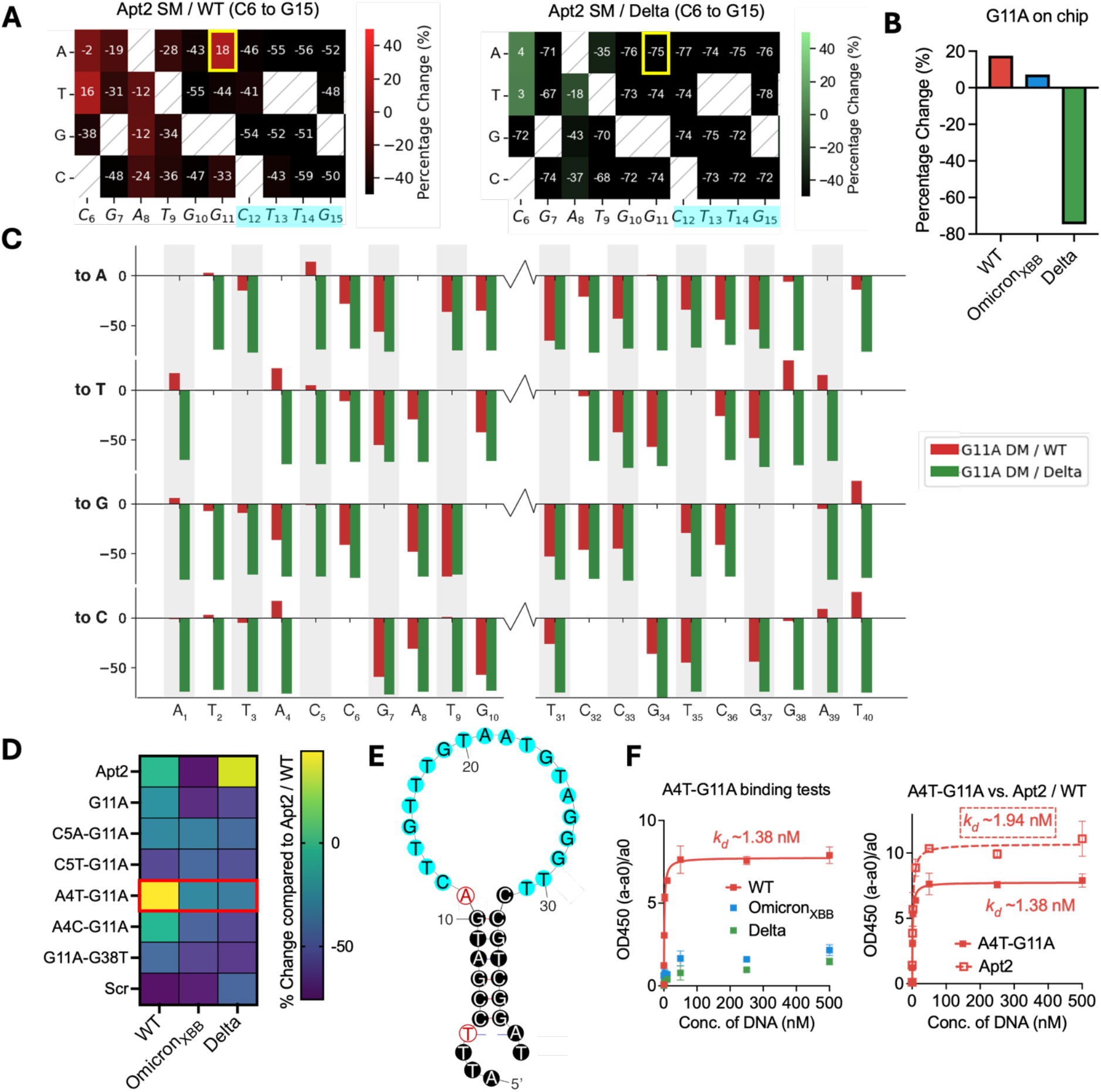
Identifying Apt2-A4T-G11A as a more selective WT binder. (**A**) Part (residues C6 to G15) of the Apt2 single-mutation/WT (left, red) and Apt2 single-mutation/Delta (right, green) heatmaps. Bases within the loop motif were highlighted in cyan. G11A point mutation cells were highlighted in yellow boxes, which exhibited that G11A single mutants (SM) sustained the binding to WT but repressed the binding to Delta. Color bars are on the right of the heatmaps. Full heatmaps can be found in **fig. S6A** and **Fig. 3A**, respectively. (**B**) According to *MiSeq* screening, G11A’s bindings to WT and Omicron_XBB_ are similar to those of the original Apt2 (+18% and +7%), but G11A has a much lower binding to Delta (-75%). **(C)** Bar charts showed selected G11A-containing DMs’ binding characteristics for WT (red) and Delta (green). The x axis represented the second mutation position; y axis showed the percentage change of the corresponding DM. (**D**) Streptavidin-mediated surface binding assay was used to test the bindings of WT-specific candidates selected from (**C**). The color scale is shown on the right. (**E**) A4T and G11A substitutions (highlighted by the red circles) were indicated in the mFold-predicted secondary structure of Apt2. The motif was highlighted in cyan. (**F**) ELONA showed that A4T-G11A remains to bind WT strongly (*k*_*d*_ ∼1±0.04 nM vs. *k*_*d*_ ∼2±0.4 nM, right), while it efficiently suppresses the binding to Delta and showing no detectable binding to Omicron_XBB_ (left).

For validation, we applied the streptavidin-mediated surface binding assay (**fig. S1B**) to a subset of G11A-containing double mutants. Mutants carrying substitutions at A1 or T40 were excluded, as they are more sensitive to be affected by interactions with chip adapters (**fig. S4**). All remaining candidates showed disrupted binding to Delta compared to Apt2. Among them, the point mutant G11A and the double mutants C5A-G11A, C5T-G11A, and G11A-C38G also exhibited noticeable reductions in WT signals (**Fig. 4D**). In contrast, the double mutants A4T-G11A and A4G-G11A displayed a WT-specific binding feature, with A4T-G11A (**Fig. 4E**) retaining the strongest binding to WT. We characterized the selected mutant A4T-G11A’s binding kinetics toward the indicated strains (**Fig. 4F**). A4T-G11A retained strong binding affinity for WT, comparable to that of the parental Apt2 (*k*_*d*_ ∼1±0.04 nM vs. 2±0.4 nM), while binding to Delta was markedly suppressed (*k*_*d*_ ∼89±10 nM vs. undetectable). These results confirmed A4T-G11A as a strong and WT-specific binder for the SARS-CoV-2 spike protein trimer, demonstrating that sequence-level screening can simultaneously enable affinity optimization and effective counterselection.

### Analyze the Binding Pattern of Apt2 with *MiSeq* Screening Data and Molecular Dynamics Simulations

Our platform can provide information beyond screening, by elucidating binding patterns useful for rational design. Initially, it was assumed that the loop motif (residues C12 to T31) was solely responsible for binding WT, with the stem serving only as a structural scaffold or a linker. However, this assumption was challenged by several observations. First, the previously identified Delta-enhanced binder A8G-A39T, which also showed improved WT binding, converted the A8·T35 Watson-Crick base pair into a G8·T35 wobble pair, a change expected to destabilize the structure. Second, the single mutant T9C, which should stabilize the stem by converting a T9·G34 wobble pair into a C9·G34 WC pair, instead completely abolished binding to all SARS-CoV-2 spike protein variants, both on chip (**fig. S9A**) and in ELONA (**Fig. 6A**). Further analysis revealed a functional asymmetry within the stem region. The upper three base pairs (T9·G34 to G11·C32) were highly intolerant to substitution by alternative WC or wobble pairs, suggesting that their role in binding extends beyond simple structural stabilization (**Fig. 6C**, above the dashed line). In contrast, the lower four base pairs (C5·G38 to A8·T35) exhibited substantially greater tolerance to base-pair alterations (**Fig. 6C**, below the dashed line), indicating a more limited contribution to protein binding.

**Fig. 6.**
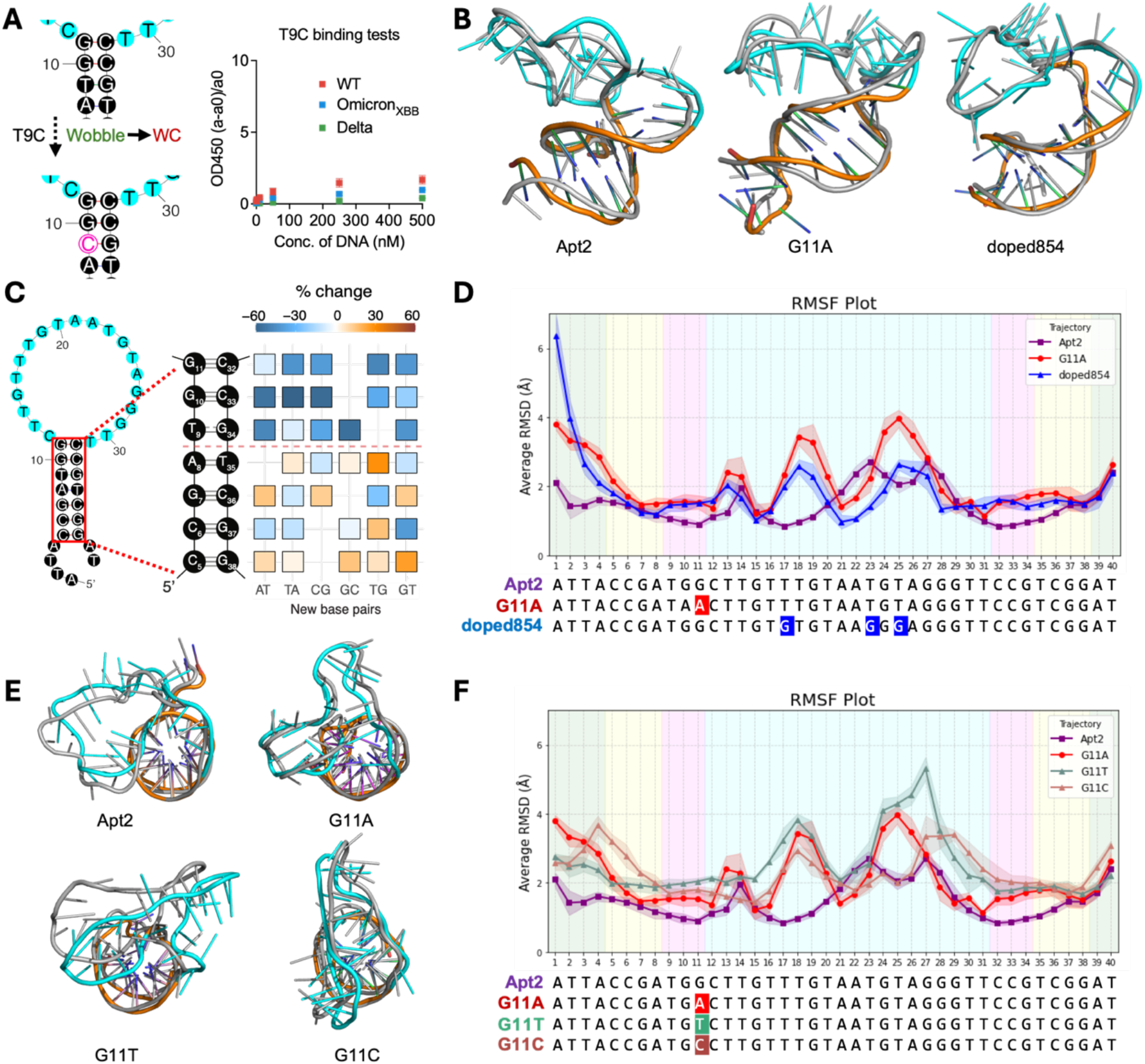
Structure analysis with on-chip data and MD simulation. (**A**) ELONA for Apt2-T9C, indicating nearly no-binding to all spike protein variants. (**B**) Representative structures in each of the top two cluster superpositions within the latter 5 ns during the 10-ns simulation time of Apt2 (left; 33.6% and 33%), G11A (middle; 28.8% and 25%), and doped854 (right; 29.8% and 29.6%). The loop motifs were highlighted in cyan. The second top structures were highlighted in grey. (**C**) Base-pair-substitution heatmap for WT screening in the stem section. The x-axis indicates the new base pair for substitution; y-axis refers to the base pair being replaced. The pink dotted line separated base pairs with higher (below) and much lower (beyond) tolerance towards base-pair substitution. (**D**) RMSF (root-mean-square fluctuation) of Apt2, G11A and doped854. (**E**) Representative structures (top views) in each of the top two cluster superpositions within the latter 5 ns during the 10-ns simulation time of Apt2 (top left; 33.6% and 33%), G11A (top right; 28.8% and 25%), G11T (bottom left; 33.2% and 22.6%) and G11C (bottom right; 37% and 30.2%). The loop motifs were highlighted in cyan. The second top structures were highlighted in grey. (**F**) RMSF of Apt2, G11A, G11T and G11C. Color code for (**D**) and (**F**): tails: green; lower stem: yellow; upper stem: magenta; loop (motif): cyan.

To provide a molecular-level insight into the possible aptamer binding pattern, we further applied molecular dynamics (MD) simulations to investigate the structural behavior of the aptamer strands (**fig. S13-14**)^76^. Structural superpositions of two most populated cluster representatives (**Fig. 6B**), RMSF (<u>R</u>oot-<u>M</u>ean-<u>S</u>quare <u>F</u>luctuation)^77^ results (**Fig. 6D**) and the simulation trajectories (step = 100, **Supplementary Movies 1-3**) showed that Apt2, G11A and doped854 had very flexible tail parts (residues A1 to A4, A39 to T40, green), and equally rigid stem (C5·G38 to G11·C32, magenta) through the simulations, suggesting that (i) truncating the 4-nt tail might not affect the binding^78^; (ii) The upper stem (T9·G34 to G11·C32) remains paired as predicted by mFold, despite their additional role beyond structural stabilization; (iii) the alteration of the binding profiles could ascribe to the different sub-structures formed within the loop. Within the loop, residues T16 to T20 in the original Apt2 showed greater stability during the 10-ns simulation compared to the Delta non-binders G11A and doped854. For the Omicron_XBB_, residues A21 to T25 displayed distinct stabilization, coinciding with two of the three mutations that form a consecutive G tract and lead to an alternative folded conformation, which aligns with the observations from the ThT experiments (**Fig. 4G**). The loop regions exhibited sharp bending towards the stems, suggesting that the role of the upper stem might be to turn the aptamer into a preferred orientation to facilitate the binding, rather than to directly contact the proteins.

In addition, according to the RMSF plot, the G11A single mutant did not pose too much effect on the behavior of itself or neighboring bases; instead, it probably altered the binding selectivity by inducing the structure change in the loop. To investigate the G11A single mutant’s binding mechanism, we performed two more simulations on G11C and G11T for comparison (**figs. S15-16**). On one hand, G11A and G11T share relatively similar conformations; however, with the latter one’s structure much more dynamic and unstable according to the RMSF analysis (**Fig. 6F**). On the other hand, G11C held a distinct 3D structure compared to G11A, G11T and the original Apt2: G11C exhibits a uniquely inward-collapsed loop architecture centered along the short stem axis, whereas Apt2, G11A and G11T adopt a more open and outward-projecting loop conformation, resulting in highly similar overall topologies which could enable the WT binding (**Fig. 6E**).

### Selective Diagnostics for One of the Three Variant Strains with Strand Displacement Reaction

Here we demonstrate an application of the optimized aptamer through our platform, by converting the highly WT-selective aptamer A4T-G11A (**Fig. 5**) into a strand-displacement fluorescent sensor^79,80^ specifically for SARS-CoV-2 WT spike protein detection. This design aimed to enhance diagnostic precision, as highly specific detection of WT enables unambiguous strain assignment, providing a foundation for generating strain-resolved data to inform studies of viral evolutionary pathways.

We have proven that the lower stem of the aptamer can be replaced by other base pairs (**Fig. 6C**), however, they cannot be truncated directly, based on the on-chip profiling results (**fig. S9C**). ELONA validation confirmed that truncation at C5·G38 yielded the shortest construct that retained strong WT binding (**fig. S19**). A4T-G11A was truncated, had the lower stem scrambled, and was extended on the 5’ end^81–84^ and labeled with FAM (denoted as wt sensor), allowing hybridization with an 11-nt Iowa Black quencher-labeled complementary strand (Q-cDNA, **Fig. 7A**). Quenching efficiency and baseline stability were evaluated prior to sensing measurements, with ∼86% fluorescence quenching achieved at a 1:2 aptamer: Q-cDNA ratio (**fig. S20A**).

**Figure 7.**
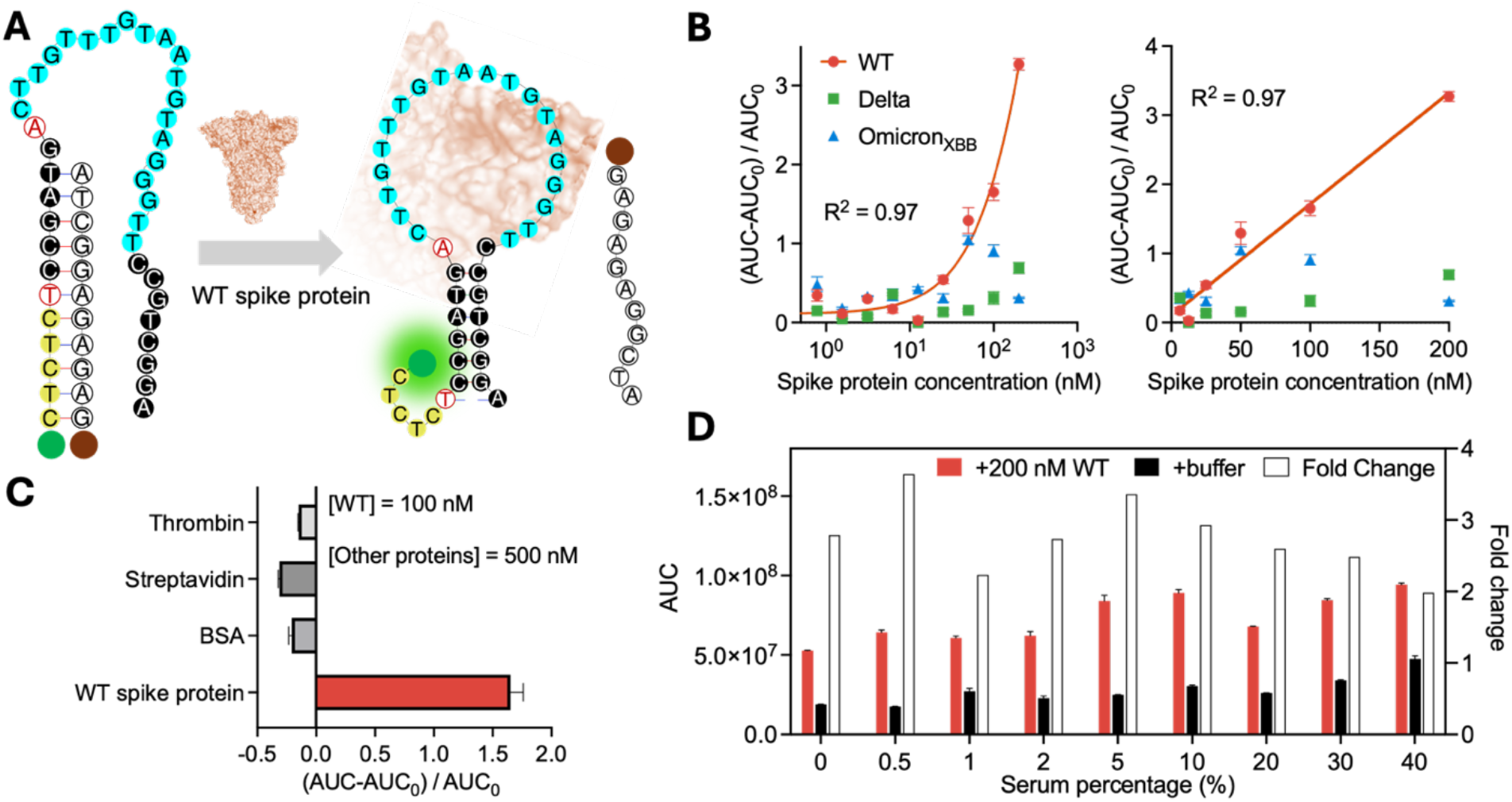
Precise diagnostics for SARS-CoV-2 WT with highly selective Apt2 mutant identified from the chip. (**A**) Schematic of the strand-displacement fluorescent sensor for rapid detection of SARS-CoV-2 spike proteins. The aptamer is modified with FAM (wt sensor), and the complementary DNA (cDNA) strand is modified with an Iowa Black quencher. Color code: loop: cyan; mutated bases: red circles; 5’ overhang: yellow. (**B**) Calibration curves for WT spike protein detection. Data were fitted using a four-parameter logistic (4PL) model over the full concentration range (0.4-200 nM, left) and a linear model over the higher concentration range (6.25-200 nM, right). The limit of detection (LOD) was determined from the linear fit. (**C**) Selectivity analysis of the aptasensor against a panel of proteins. Non-target proteins were tested at 5× higher concentrations than the WT spike protein. (**D**) Stability and reactivity of the WT selective aptasensor in response to 200 nM WT spike protein across varying serum percentages (0-40%). For (**B**)-(**D**), each condition was measured in duplicate.

Upon addition of 200 nM WT spike protein, fluorescence was enhanced by ∼3 fold, significantly exceeding the responses observed for Delta and Omicron_XBB_ at the same concentration (**fig. S20B**). A calibration curve was constructed from the fluorescence enhancement upon WT addition with a four-parameter logistic (4PL) model (**Fig. 7B, left**)^85^. The response exhibited a linear relationship above 6.25 nM (**Fig. 7B, right**), from which a limit of detection (LOD) of 5.5 nM was estimated using 3σ/slope. In contrast, the responses to Delta and Omicron_XBB_ remained near baseline, demonstrating strong specificity for the WT strain. We further evaluated sensor selectivity against a panel of common proteins (**Fig. 7C**). All non-target proteins produced negligible fluorescence changes, even at 5-fold higher concentrations (500 nM). Finally, the aptasensor maintained robust performance in complex biological matrices, retaining ∼2-fold signal enhancement in 40% human serum (**Fig. 7D**), supporting its potential applicability in clinical samples^86–88^.

## DISCUSSION

Here we demonstrate a massively parallel strategy for rapidly diversifying and screening spike-protein-binding aptamers within days, enabling either retention of binding to newly emerged viral strains or enhancement of specificity toward a particular strain. Starting from an early WT-spike protein aptamer Apt2, we were able to identify a mutant with improved binding to the Delta strain, a nM-level binder for Omicron_XBB_, and a WT-specific mutant with the binding selectivity significantly enhanced. The platform also provided systematic exploration of the sequence-function relationship, revealing the binding pattern structurally incorporated with molecular dynamics simulations. Finally, we turned the selected WT-specific aptamer into a strand-displacement fluorescence sensor, enabling the precise diagnostics for SARS-CoV-2 WT.

Although virus-targeting aptamers are unlikely to replace monoclonal antibodies (mAbs) in the foreseeable future because of intrinsic limitations, including rapid renal clearance, susceptibility to nuclease degradation, and limited ability to engage immune effector functions^89–91^, nevertheless, aptamers remain attractive for biosensing and certain therapeutic applications because of their low production cost, high stability, small size, and low immunogenicity^92–94^. Our results highlight an additional advantage of aptamers: adaptability. Once an initial antiviral aptamer has been identified, it can be rapidly reengineered to recognize newly emerging variants without requiring de novo selection each time. In addition to improving binding to evolved strains, this study also demonstrates that aptamers can be tuned for strain-selective recognition, which may be particularly valuable for diagnostic applications requiring precise variant discrimination and for generating strain-resolved datasets to support studies of viral evolution^75,95–97^. We also demonstrated that aptamers offer functionality beyond binding^98^ by converting a WT-selective variant into a strand-displacement fluorescent sensor. Their ability to be readily engineered into structure-switching aptasensors^79,88^ provides a functional advantage over mAbs in certain sensing and diagnostic applications.

Repurposed NGS platforms offer a unique advantage for aptamer engineering by enabling massively parallel and quantitative interrogation of large, closely related sequence spaces in a single experiment. Prior studies^36,40,43–45,47,48,53,59^, including our own^55,57,58,99^, have demonstrated the versatility of this approach across diverse molecular systems, establishing repurposed NGS flow cells as a broadly useful platform for functional screening and sequence-property mapping. In this context, the key strength of NGS-based screening is not only throughput, but also the ability to retain and evaluate sequence diversity directly, rather than relying on iterative enrichment steps. This work extends this capability to a distinct challenge of rapid adaptation of antiviral aptamers to continuous viral evolution. Rather than treating each novel viral variant as a de novo discovery problem, we show that a repurposed NGS platform can be used to reengineer an existing functional aptamer through a random-rational hybrid diversification around a validated seeding aptamer. This strategy preserves the efficiency of high-throughput screening while introducing a more directed and adaptable framework for molecular redesign. As a result, the platform supports both rapid recovery of binding toward newly emerged variants and deliberate tuning of strain specificity, highlighting a specific advantage of our approach over existing repurposed NGS applications: it is not limited to characterizing or optimizing static molecular functions, but can instead be used to keep pace with an evolving biological target. The ability to iteratively screen related sequence libraries on the same platform further strengthens its practical value for emerging-pathogen surveillance and response.

Beyond the *MiSeq* screening, molecular dynamics simulations provide a structural framework for interpreting the variant-dependent binding behaviors observed experimentally (**Fig. 6**). These results suggest that the upper stem region, rather than serving solely as a passive scaffold, contributes directly to target recognition. Within the loop, distinct segments appear to govern variant-specific interactions, indicating that binding specificity is distributed across multiple localized motifs rather than confined to a single region. Notably, substitutions at position G11 exert a pronounced effect on global conformation. While the adenine substitution preserves a loop orientation compatible with binding, alternative substitutions lead to structural rearrangements or destabilization that disrupt target interaction. This provides a structural explanation for the experimentally observed selectivity at this position. Importantly, this suggests that aptamer optimization need not rely solely on empirical screening, but can be guided by structural considerations, providing a foundation for rational aptamer design in response to evolving viral targets.

More broadly, this work addresses a challenge highlighted by the COVID-19 pandemic: the lack of a modular molecular-recognition platform that can be readily reprogrammed to keep pace with viral evolution. Although the demonstration here was SARS-CoV-2 spike protein, this strategy should be extendable to other viral targets. In a future outbreak scenario, this could enable a practical workflow in which libraries diversified around an existing antiviral aptamer are prepared on repurposed NGS flow cells and screened immediately against newly emerging variants.

The library size can also be scaled up by utilizing other types of NGS flow cells (*MiSeq, HiSeq* and *NextSeq* and *NovaSeq*) or leveraging multiple chips. Because these chips can remain functional for years under appropriate storage (when soaked in an appropriate buffer and stored at 4 °C) and can be reused for sequential assays, the same platform could support iterative screening as new strains arise, without requiring complete library redesign unless full target escape occurs.

In summary, we developed a new method to rapidly evolve aptamers against newly emerging viral strains, providing a potential tool for responding to current and future pandemics. We demonstrated its effectiveness by identifying sequences that bind the 9-amino-acid-mutated Delta variant and the 39-amino-acid-mutated Omicron_XBB_ variant within a diversified pool based on Apt2, which was originally selected against the baseline strain WT. This streamlined workflow can be readily adapted to a broad range of biomedical applications in the future.

## MATERIAL AND METHODS

### Doped pool mutation probability calculation

Let the number of the doped bases be L and the mutation rate be m. The theoretical number of the mutated bases n follows the binomial distribution.

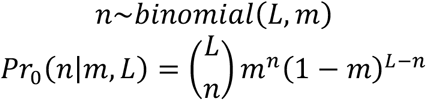

With the n>2 filtration, the ultimate probability of n given m and L can be calculated as follows.

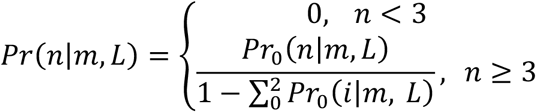

In this study, we set L = 20 (only dope the loop region) and m = 0.1. The theoretical and actual frequency of each mutation number can be found in **fig. S2**.

### Apt2 library construction for NGS sequencing

Apt2 variants library with PCR primers was synthesized by Twist Biosciences. Forward and reverse PCR primers with Illumina sequencing adapters were synthesized by IDT. The overlap extension PCR process was performed using Q5 High-Fidelity 2× Master Mix from NEB (#M0492) following protocol provided by the manufacturer using a thermal cycler (Applied Biosystems). After PCR amplification, the library amplicons were purified with the PCR cleanup kit (#T1030, Monarch). The length of the amplicons was checked with a 2% agarose gel (**fig. S21**). The concentration and quality of the purified library amplicons was quantified using a NanoDrop 2000 UV-Vis spectrophotometer (Thermo Scientific). After PCR process, PhiX control V3 sequences (#FC-110-3001, Illumina) were diluted and mixed with Apt2 amplicons to prepare a 20 µl of 2 nM solution according to the requirement of our sequencing facility. The coverage for both PhiX and the library sequences were targeted to be 15% ∼ 20% of overall sequencing space. The mixture was then subjected to *MiSeq* PE75 sequencing and the final loading concentration was at 8 pM.

### Illumina *MiSeq* chip preparation

After sequencing, *MiSeq* chips were stored at 4°C in 20 mM sodium phosphate buffer (SPB, pH 6.6). Before experiments, the *MiSeq* chip was incubated with 20 µl 0.1 N NaOH solution for 5 min at room temperature and rinsed with 20 µl 20 mM SPB (pH 6.8) 3 times to remove excess NaOH. This step was to denature DNA strands which might contain residual fluorescent dyes from the sequencing process. The chip was further photobleached by sequentially exposing each FOV (5 rows, 65 FOVs per row, 2 layers, 220 × 220 µm^2^ each FOV) under an 80% excitation power for 1 min. The PhiX sequences were then labeled using an Atto488-tagged probe (1 µM and 20 µl) at 37°C for 40 min. The flow cell was rinsed 3 times by the screening buffer (40 mM Tris-HCl (pH 7.5), 150 mM NaCl, 6 mM MgCl_2_, 2.5 mM CaCl_2_, 2.5 mM KCl, 0.1% Tween 20, 0.2 mg/mL BSA) to remove excess fluorophores. 20 µL 100 nM of His-tagged SARS-CoV-2 spike protein variants (#SPN-C52H3, #SPN-C52He, #SPN-C5248; Acro Biosystems) were then introduced to the chip surface by pipetting and incubated at 37°C for 1 hour. The protein was stored in nuclease-free water at 1 µM and further diluted in screening buffer at room temperature for 30 min. The flow cell was rinsed 3 times by the screening buffer to remove unbound proteins. 20 µL of 6×-His-tag mAb-Alexa647 conjugate (#MA1-21315-A647, Invitrogen) were then introduced to the chip surface by pipetting and incubated at room temperature for 15 min. The mAb-dye conjugate was diluted at 1:500 ratio with the screening buffer. Excess conjugate was washed away with screening buffer.

### Fluorescence microscopy and chip image acquisition

An open-source software, Micro-Manager^100^, was used to control an sCMOS camera (ORCA-Flash 4.0, Hamamatsu), an xyz translation stage (ProScan III, Prior Scientific), and an auto-shutter (Lambda SC, Shutter Instrument) on an Olympus IX71 fluorescence microscope for all our *MiSeq* screening experiments. An LED illuminator and a 60× water-immersion objective (UP-LSAPO60XW, Olympus) were used in the IX71 system. The detailed setup of images acquisition could be found in our previous reports^55,57^. In brief, for each round of experiments, we acquired 3 rows of TIFF images with 60 field of views (FOVs, 220×220 µm^2^) per row on both sides of the chip (totally 360 images). This covers 5.81 mm^2^ surface area. For each row of images, we first recorded fiducial marker images (1 second exposure time, green channel, Ex/Em: 480/40 nm, 535/50 nm), and then acquired spike protein images with red channels (Ex/Em: 620/60 nm, 700/75 nm, cat. no. 49006, Chroma) under the same imaging settings. The scanning step size (220 µm per step) and exposure time at each position were all precisely controlled by our auto-scan algorithm and shutter control.

### Illumina *MiSeq* chip regeneration

After loading the samples to the chip and acquiring the images, the flow cell was regenerated by digesting the proteins and denaturing the anchored DNA sequences. 20 µL proteinase K mix (#P8107, NEB) was added to the flow cell by pipetting and incubated at 40°C for 2 hours. Excess proteinase and protein fragments were washed away by 20 mM SPB buffer. This was followed by a 0.1 N NaOH denaturation. Excess base was removed by 20 mM SPB buffer.

### Fluorescence image alignment algorithm

To identify the sequences behind polonies, we applied a revised imaging alignment algorithm from <u>**C**</u>hip-<u>**H**</u>ybridized <u>**A**</u>ssociated <u>**M**</u>apping <u>**P**</u>latform, or CHAMP ^53^, which was developed by Jung *et al*. The detailed description of implementing CHAMP algorithm on epi-fluorescence-microscope-acquired images could be found in our previous report^55,57^. In brief, FASTQ coordinates were first Gaussian blurred to build a synthetic image and then transformed into Fourier space. TIFF images were also Fourier transformed and performed cross-correlation with transformed FASTQ images to register the rough position of each TIFF image. If the maximum cross-correlation value is larger than that of control position (other positions with higher read density), this TIFF image will perform precision alignment using least-squares constellation mapping to the potential target region to find detailed parameters (e.g., scaling factor, rotation angle). The output files included the x-y coordinates and the corresponding sequence ID of each cluster within the FOV and were saved individually according to different image positions. To analyze our Apt2 images, we corrected the uneven illumination using flat-field correction and bootstrapped to compute the median intensity of each activator (baseline corrected) in order to rank the Apt2 mutants’ brightness.

### Flat-field correction

To correct the uneven illumination across each FOV. We implemented flat-field correction. A Gaussian-blur filter (σ = 100) was applied on each FOV itself to generate the flat-field reference image for each FOV. The corrected images used for the following analysis were computed as follows:

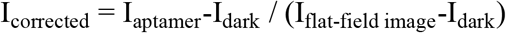

Where I_corrected_ is the flat-field-corrected images, I_aptamer_ is the original TIFF images acquired under red channel, I_flat-field image_ is the TIFF image after Gaussian blurring, and I_dark_ is the dark image acquired using 1 second exposure time when illumination source is turned off. When calculating the percentage changes, the intensities were subtracted by 1 to remove the average background from the protein and dye’s non-specific binding to the chip surface.

### ELONA binding assay

The binding affinities quantified in this study were all produced with ELONA ^63^ (enzyme-linked oligonucleotide assay). Briefly, 96-well high-binding microplates (#3690, Corning) were coated with the SARS-CoV-2 spike protein variant at a concentration of 1.33 µg/mL in PBS (pH 7.4, Invitrogen #AM9624) and incubated at 4 °C overnight. After washing with PBST (PBS containing 0.05% Tween-20), wells were blocked with blocking buffer (5% w/v BSA in PBST) for 1 hour at room temperature. Biotin-labeled aptamers (IDT) at various concentrations (3 duplicates for each concentration) were then added and incubated for 90 min at room temperature to allow binding. After washing, bound aptamers were detected using streptavidin-HRP conjugate (#RABHRP3-600UL, Millipore Sigma), followed by addition of TMB substrate (#555214, BD Bioscience). Washing was introduced between the steps with the screening buffer. The reaction was stopped with stop solution (10% H_2_SO_4_), and absorbance was measured at 450 nm using a microplate reader (SpectraMax i3, Molecular Devices). Control with scrambled Apt2 (Scr) were included. OD450 values were normalized with the lowest value on the whole plate.

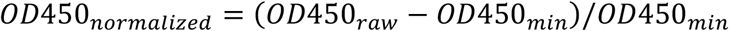

The dissociation constant (*k*_*d*_) was generated by fitting the concentration (c)-absorbance (A) curve to a Hill equation with a dynamic Hill slope (h), given that the signal produced by an enzymatic reaction might not be in a strictly linear relation with the binding. Each of the 3 duplicates was fitted to the model separately. The reported *k*_*d*_s are composed of the mean *k*_*d*_ of the duplicates with the standard deviation:

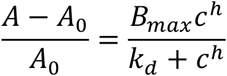

### Streptavidin-mediated surface binding assay

In addition to ELONA, streptavidin-mediated surface binding assay was also used to estimate and compare the bindings for the DNA sequences and spike protein variants. Biotin-labeled aptamers (IDT) at 2.5 µM (3 duplicates) were then added to the 96-well streptavidin-coated pre-blocked black microplates (#15503, Thermo Fisher). and incubated for 2 hours at room temperature to be anchored onto the plate surface. After washing, 33 nM spike protein variants were added and incubated for 1 hour to allow binding, followed by addition of the 0.002× 6×-His-tag mAb-Alexa488 conjugate (#MA1-21315-A488, Invitrogen) for 30 min at room temperature. After washing away excess conjugate, 100 µL screening buffer was added to each well to enable the fluorescence measurement. Fluorescence was measured at Ex/Em 485/535 nm using the microplate reader. Control with scrambled Apt2 (Scr) were included.

#### 3D structures prediction for Apt2 and its mutants of interests

Starting from the predicted secondary structure of Apt2 by mfold^62^, we replaced all Ts with Us to generate its equivalent ssRNA rApt2. The corresponding 3D structures rApt2 was then modeled and visualized in RNAComposer^101,102^. The preliminary 3D structure of Apt2 was then obtained by substituting bases T back for U. To relax the aptamer system, a refinement process was carried out based on molecular dynamics simulation to produce the input structure for the MD simulation. The input structures of G11A, doped854, G11C, G11T was generated by performing mutagenesis to the Apt2’s 3D structure by PyMol^103^.

### Molecular dynamics simulation of selected aptamers

The 3D structures of Apt2 and mutants of interests were obtained as described in the previous section. The structures were then soaked into a box of water with a size of 65Å × 65Å × 65Å. Counter ions such as Na^+^ were added to neutralize the system. Additional NaCl, KCl, MgCl_2_ and CaCl_2_ were added to obtain salt concentrations of 150, 6, 2.5 and 2.5 mmol/L, respectively, to mimic the correct buffer condition. The systems underwent restrained minimization and restrained gradual heating processes to reach equilibration before 10-ns production simulation for analysis^104,105^. All simulations in this study were performed with the AMOEBA polarizable force field^106–108^.

### Simulated frame clustering

A pairwise RMSD (Root mean square deviation) matrix was computed from the aligned frames to serve as the distance metric. We applied agglomerative hierarchical clustering to this matrix, iteratively merging the closest clusters based on a complete linkage method. The resulting dendrogram was cut at a threshold of RMSD = 4 to partition the trajectory into distinct conformational states. and then the centroid is taken as representative structure. The central frames were collected from each cluster and superposed to visualize for each mutant of interests.

### RMSF calculations

Root mean square fluctuation (RMSF) analysis was conducted to evaluate the flexibility of individual residues within the backbones of aptamer structures during MD simulations. The RMSF values were calculated over the equilibrated trajectory using PyLOOS, according to the equation:

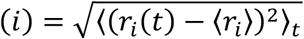

where r_i_(t) represents the position of atom i at time t, and ⟨r_i_⟩ denotes its time-averaged position after aligning each frame to the first one. Only the equilibrated portion of the trajectory (last 5 ns) was used for the calculation.

### Strand-displacement fluorescent sensor test

The fluorescence aptasensor experiments were performed in a half-area black 96-well microplate (#675076, Greiner Bio-One) using the microplate reader mentioned in the previous section. The excitation wavelengths were set to 480 nm; the emission spectra were collected from 510-700 nm. 2.5 µL of 1 µM wt sensor, 40.5 µL of screening buffer and 2 µL of proteins at different concentrations were premixed in the wells and incubated at RT for 30 min. 5 µL of 1 µM Q-cDNA was then added the wells before measuring the fluorescence. The fluorescence intensity for each sample was calculated through the area under curve (AUC) of the emission spectrum. For the spike-in serum test, 50% human serum was used to yield buffer with 0-40% serum. The mixture was heat-inactivated at 85°C for 10 min. The supernatant was collected as the new buffer. The data were represented as (AUC-AUC_0_) / AUC_0_, where AUC_0_ denotes the fluorescence with the mixture of 2 µL of screening buffer, and AUC was the fluorescence with 2 µL of protein addition.

## ACKNOLEDGEMENTS

We thank L. Xu for helping generate **Fig. 5C** and **Fig. 6C**.

## Author Contributions

Y. H., Z. Y., Y.-A. K., Y. Wu, Y. L. and H.-C. Y. discussed and defined the project. H.-C. Y. and Y. L. supervised the project. Y. H. and Y.-A. K. designed the screening library. Y. H. and K.K. L. and W.-R. C. performed chip screening experiments. Y. H. and Y.-A. K. analyzed the champ results. Z. Y., Y. Wu and A.-T. N. helped design the binding characterization assays. Y. H. and D. F.-A. performed the binding characterization assays. J. G., Y. Wang and Y. H. performed and analyzed MD simulations. Z. Y. and Y. H. designed and performed the strand-displacement fluorescent sensor tests. Y. H. and Y. C. developed Python and R scripts for data illustration. Y. H. and K.K. L. performed gel electrophoresis experiments during the optimization phase. Y.-I. C., S. H., T.D. N., S. K. and P. R. provided critical insights and contributed to the discussion. S. S. optimized the buffer condition. Y. Wu, Y.-A. K., Z. Y., Y.-I. C and D. F.-A. reviewed or revised the manuscript. Y. H. and H.-C. Y. wrote the article with editorial assistance from all co-authors. H.-C. Y. supervised the project.

## Data and materials availability

All data needed to evaluate the conclusions in the paper are present in the paper and/or the Supplementary Materials. Other data are available from the corresponding author upon reasonable request.

## Supplementary Materials

Tables S1 to S3, Notes, Supplementary Figs. S1 to S21, Movies S1 to S3

