## Supplementary Materials for "A High-Throughput Platform for Rapid Adaptation of DNA Aptamers to SARS-CoV-2 Evolution"

**Table S1. Sequences used in this study**

For the predicted stem-loop structures, loop regions are denoted by underscores. Mutated bases are colored green, cyan, red, and yellow for Delta-, Omicron$_{XBB}$-, WT-candidate binders, and other strands, respectively. The partially hybridized regions in SP1 and SP2 are indicated in bold.

| Name | Sequence |
|---|---|
| Aptamers reported by other groups mentioned in this study | |
| Apt2 [1] | ATTACCGATGGCTTGTTTGTAATGTAGGGTTCCGTCGGAT |
| MSA1-T2 [2] | TTCCGGTTAATTTATGCTCTACCCGTCCACCTACCGGAA |
| AR10 [3] | CCCGACCAGCCACCATCAGCAACTCTTCCGCGTCCATCCCTGCTG |
| CoV2-RBD-1C [4] | CAGCACCGACCTTGTGCTTTGGGAGTGCTGGTCCAAGGGCGTTAATGGACA |
| Apt1 [1] | TCGAGTGGCTTGTTTGTAATGTAGGGTTCCGGTCGTGGGT |
| CoV2-6C3 [5] | CGCAGCACCCAAGAACAAGGACTGCTTAGGATTGCGATAGGTTCGG |
| MSA1-T6 [2] | TTACGTCAAGGCTTTCCGGTACCGGAAGCATCTCTTTGGCGTG |
| Mutants in this study | |
| T3C-A39C | ATCACCGATGGCTTGTTTGTAATGTAGGGTTCCGTCGGCT |
| ins3C | ATCTACCGATGGCTTGTTTGTAATGTAGGGTTCCGTCGGAT |
| A8G_A39T | ATTACCGGTGGCTTGTTTGTAATGTAGGGTTCCGTCGGTT |
| C36T_T40A | ATTACCGATGGCTTGTTTGTAATGTAGGGTTCCGTTGGAA |
| T3G_G38C | ATGACCGATGGCTTGTTTGTAATGTAGGGTTCCGTCGCAT |
| G19T-G29T | ATTACCGATGGCTTGTTTTTAATGTAGGTTTCCGTCGGAT |
| doped3557 | ATTACCGATGGCTCGTTTAGAATGTCTGGTTCCGTCGGAT |
| T17G-G24C | ATTACCGATGGCTTGTGTGTAATCTAGGGTTCCGTCGGAT |
| doped854 | ATTACCGATGGCTTGTGTGTAAGGGAGGGTTCCGTCGGAT |
| A21T-T30C | ATTACCGATGGCTTGTTTGTTATGTAGGGCTCCGTCGGAT |
| G11A | ATTACCGATGACTTGTTTGTAATGTAGGGTTCCGTCGGAT |
| C5A-G11A | ATTAACGATGACTTGTTTGTAATGTAGGGTTCCGTCGGAT |
| C5T-G11A | ATTATCGATGACTTGTTTGTAATGTAGGGTTCCGTCGGAT |
| A4T-G11A | ATTTCCGATGACTTGTTTGTAATGTAGGGTTCCGTCGGAT |
| A4C-G11A | ATTCCCGATGACTTGTTTGTAATGTAGGGTTCCGTCGGAT |
| G11A-G38T | ATTACCGATGACTTGTTTGTAATGTAGGGTTCCGTCGTAT |
| T9C | ATTACCGACGGCTTGTTTGTAATGTAGGGTTCCGTCGGAT |
| G4Apt2 | ATTACCGATGGGTTGGGTGTAAGGGAGGGTTCCGTCGGAT |
| *Miseq* chip-related sequence | |
| P5 | AATGATACGGCGACCACCGAGA |
| P7 | ATCTCGTATGCCGTCTTCTGCTTG |
| SP1 | TCTACACTCTTTCCCTACACGAC**GCTCTTCCGATCT** |
| SP2 | **AGATCGGAAGAGC**ACACGTCTGAACTCCAGTCAC |
| Barcode | ATCACG |
| PhiX probe | /5ATTO488N/TTTCGGTCTCGGCATTCCTGCTGAACCGCTCTTCCGATC |

| | |
|---|---|
| FP | AATGATACGGCGACCACCGAGATCTACACTCTTTCCCTACACGACGCTCTTCCGATC |
| RP | CAAGCAGAAGACGGCATACGAGATCGTGATGTGACTGGAGTTCAGACGTGTGCTCTTCCGATC |
| Sensor | |
| wt sensor | /56-FAM/CTCTCTCGCTTGACTTGTTTGTAATGTAGGGTTCCGAGCGA |
| Q-cDNA | AAGCGAGAGAG/3IABkFQ/ |
| wt T1 | /56-FAM/TCGCTTGACTTGTTTGTAATGTAGGGTTCCGAGCGA |
| wt T2 | /56-FAM/GCTTGACTTGTTTGTAATGTAGGGTTCCGAGC |
| wt T3 | /56-FAM/CTTGACTTGTTTGTAATGTAGGGTTCCGAG |
| wt T4 | /56-FAM/TTGACTTGTTTGTAATGTAGGGTTCCGA |
| Other | |
| Scr | TTAGGTCTTGCGGTGGCCTATTATTGTTGAATTCGAGGCA |
| cTBA | CCAACCACACCAACC |
| D-22AG-40 | AGGGTTAGGGTTAGGGTTAGGGAGGGTTAGGGTTAGGGTT |

**Table S2. Amino acids mutated in the SARS-CoV-2 variants used in this study**

| Delta [6,7] | Omicron$_{XBB}$ [7,8] | | | |
|---|---|---|---|---|
| T19R | T19I | G339H | N440K | Y505H |
| G142D | L24S | R346T | V445P | D614G |
| E156G | Del25/27 | L368I | G446S | H665Y |
| Del157/158 | V83A | S371F | N460K | N679K |
| L452R | G142D | S373P | S477N | P681H |
| T748K | Del144/145 | S375F | T478K | N764K |
| D614G | H146Q | T376A | E484A | D796Y |
| P681R | Q183E | D405N | F490S | Q954H |
| D950N | V213E | R408S | Q498R | N969K |
| | G252V | K417N | N501Y | |

**Table S3. Number of duplicates in the MiSeq screening experiments**

This table is specific for the seeding aptamer Apt2 and the scrambled sequence (Scr). Results are shown in **Fig. 2D-E**.

| | WT | Delta | Omicron$_{XBB}$ | Total reads from NGS |
|---|---|---|---|---|
| Apt2 | 16,238 | 9,459 | 21,163 | 35,859 |
| Scr | 26,106 | 15,152 | 34,013 | 53,023 |

### Notes

Before trying Apt2[1], we also tried a few more seeding aptamers for the library, but not all of them are compatible with our current system, even though they were characterized as strong binders. These aptamers include MSA1-T2 by Li group[9] and AR10 by Lu group[3] (**fig. S17**). We hypothesized that the incompatibility between the binding pattern and the orientations of the molecules on the flow cell might be the main cause. In addition, the flow cell has a limited lifetime due to the repeated NaOH washes and some protein debris would be stuck on the surface despite the treatment of the proteinase K, which is a source of inaccuracy.

The surface-binding assay in this study was designed to roughly estimate the binding profile of a strand in an easier and low-cost way. It mimicked the interactions taking place on the *MiSeq* chip surface, while with lower surface tension given the much larger volumes, which may allow the molecular interaction to be more closely resemble those occurring freely in solution and provide a more physiologically relevant context for the reactions. In addition, the chip-related adapters were not included in the assay, eliminating the effect of those sequences. Interestingly, from the outcome (**Fig. 4D, Fig. 5C, and Supplementary Fig. S6**), we observed that Apt2/Delta showed a stronger fluorescence signal than Apt2/WT, which was contradictory to the previous ELONA results (**Fig. 2B**). We hypothesized that the Delta spike protein trimer may adopt a conformation that more effectively exposes its C-terminal His-tags, thereby enhancing accessibility for fluorescent antibody labeling, which was supported by the crystal structures of WT and Delta trimers.

Neither the original Apt2 nor doped854 has a perfect G-quadruplex-forming pattern, while both enhanced thioflavin T assay (ThT)'s fluorescence to some extent (**Fig. 4G**). To further demonstrate that the 53% increasement from Apt2 to doped854 came from a meta-stable G-quadruplex instead of a false positive signal from a G-rich sequence, we also performed the assay on two more mutants: (i) a rational-designed G4-formed doped854 mutant G4Apt2 (C12G-T16G-T17G-T23G-T25G), which has a perfect G-quadruplex-forming pattern $G_{\geq3}N_xG_{\geq3}N_xG_{\geq3}N_xG_{\geq3}$ to ensure the activation of ThT (**Fig. 4F**); (ii) A4T-G11A, our WT-specific mutant, which has one fewer guanine than the original Apt2. G4Apt2 turned out to have a relatively stronger signal than doped854, suggesting a more stable G4, while A4T-G11A exhibited a very similar outcome to that of Apt2 (**fig. S18**), implying that the fluorescence signal was not solely dependent on the enriched guanines. Overall, we presumed that the doped854 can form a meta-stable G-quadruplex of which the original Apt2 was not capable, which may be associated with its unique $\text{Omicron}_{\text{XBB}}$-binding property.

Like many other viral S protein, SARS-CoV-2 S protein also undergoes glycosylation in the host cells[10]. The N-glycans on the proteins are believed to stabilize the structure through shielding and facilitate the ACE2 recognition by expose the RBD[11]. While there is no evidence that the glycans would have a huge impact on the S protein structures or significantly influence its functions. We used commercially available S proteins in this study, which were recombinant proteins expressed in mammalian cells. Although glycosylation they underwent might not be the same with that during actual infection events, we believe the effects would be neglectable, since recombinant S proteins by mammalian cells have been widely used in vaccine[12] and antibody[13] studies.

**Supplementary Figure 1**

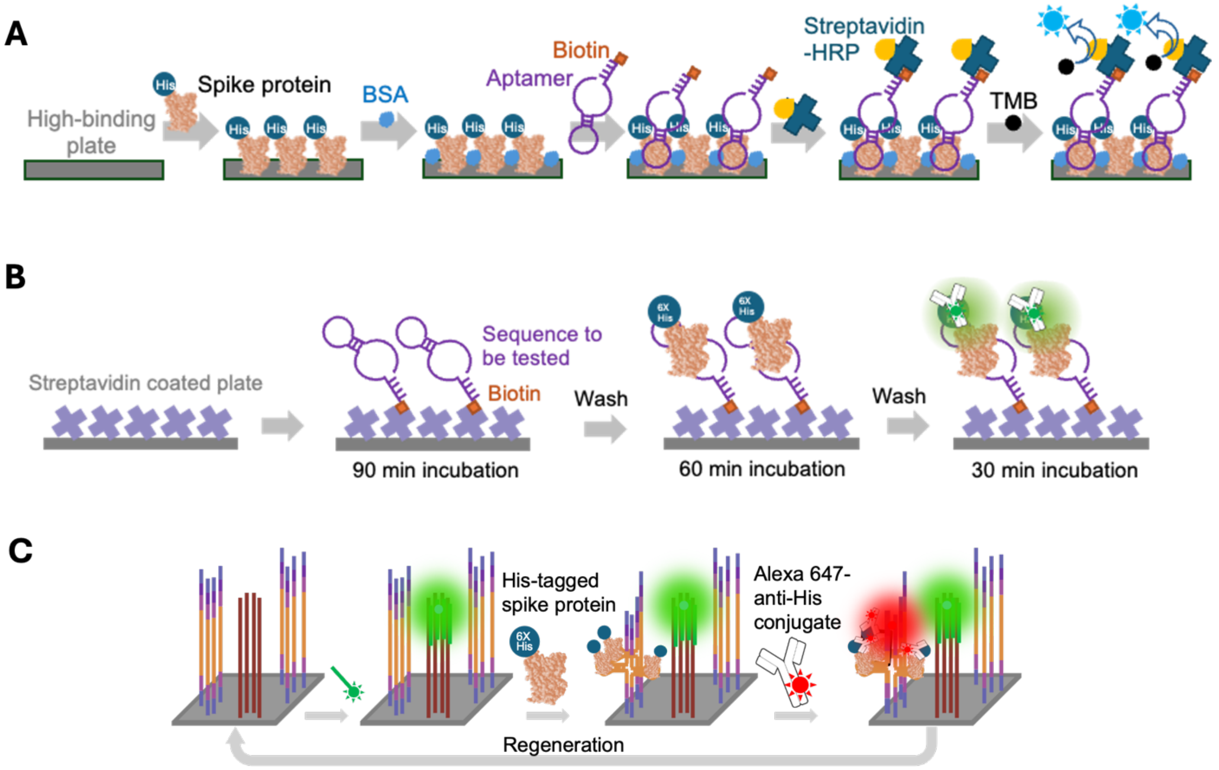

**Fig. S1. Workflows of all three binding assays employed in this study.** (**A**) ELONA (Enzyme-linked Oligonucleotide Assay) was used to generate aptamer's $k_d$ values in this study. (**B**) streptavidin-mediated surface binding assay was used to roughly estimate a specific strand's binding capability with a target protein variant. (**C**) *MiSeq* screening workflow: to image the spike protein binding on the chip, we first incubated the flow cell with 100 nM His-tagged spike protein trimer, which was then stained by the Alexa647 His-tag antibody conjugate. After imaging, the chip was regenerated by 0.1 N NaOH treatment, with another spike protein variant added in the following round. Refer to **Fig. 2** and **Methods** for more details.

**Supplementary Figure 2**

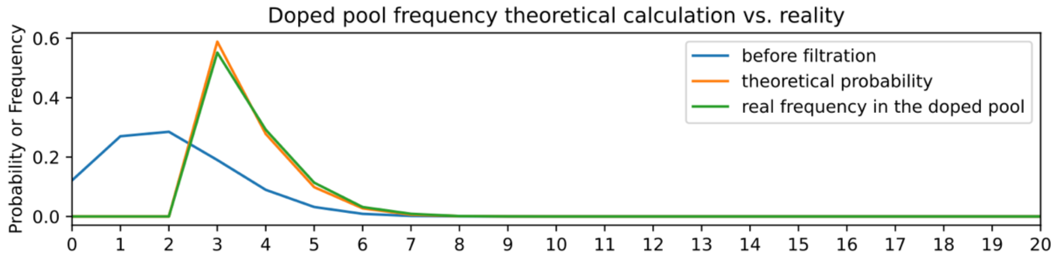

**Fig. S2. Doped mutant generation.** The doped mutants were generated with a mutation rate of 10% [14] within the 20-nt loop motif (binomial (20, 0.1), blue) and filtered to remove duplicates with the other subgroups (yellow). The mutated base numbers of final 4,500 doped sequences were calculated and shown in green, which was well aligned with the theoretical probability.

**Supplementary Figure 3**

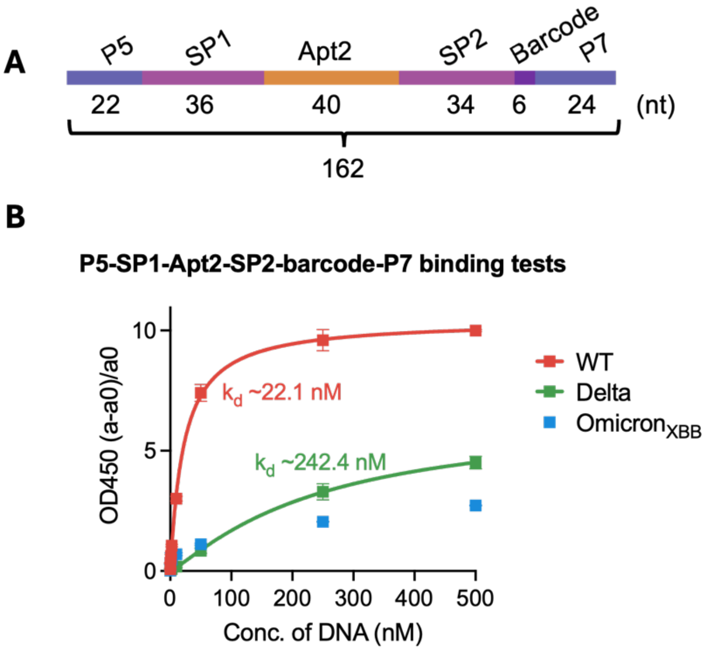

**Fig. S3. The adaptors on the chip would not completely inhibit Apt2's bindings.** (**A**) The component of the tested sequence, which is exactly how the aptamer pool would be presented on the Illumina *MiSeq* flow cell. The bottom row indicates the length of each part. The whole strand was 162 nt long. The exact sequence can be found in **Supplementary Table S1**. (**B**) Use ELONA to quantify the binding affinities of the sequence described in **fig. S1A**, with $k_d$ (WT) ~22 nM and $k_d$ (Delta) ~242 nM. Each sample has 3 duplicates. Error bars represent the standard deviations.

### Supplementary Figure 4

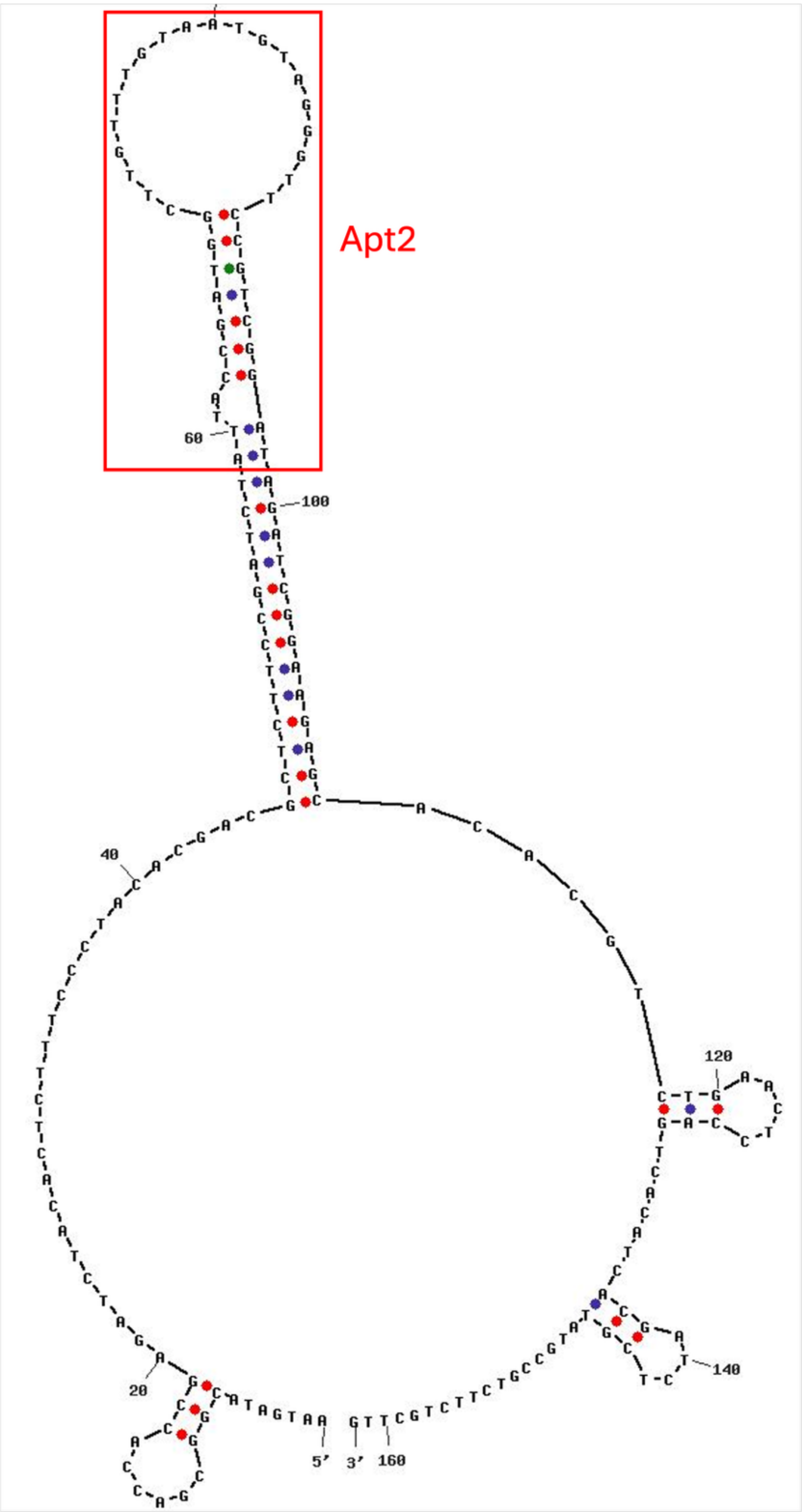

**Fig. S4.** Secondary structure of P5-SP1-Apt2-SP2-barcode-P7, predicted by UNAFold [15]. Apt2 sequences [1] was highlighted in red.

### Supplementary Figure 5

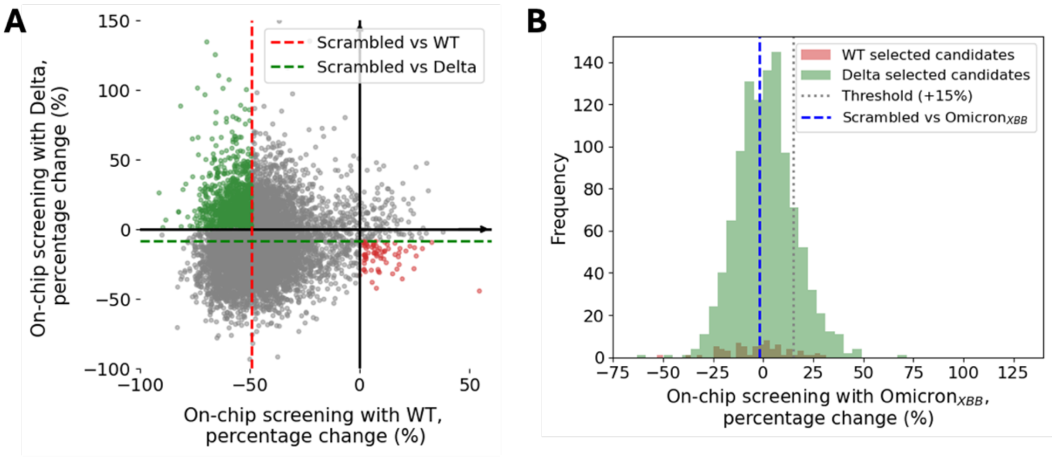

**Fig. S5. Combinatorial analysis for WT and Delta data.** Sequences with intensities higher than Apt2/WT and lower than Scr/Delta (red circles in **A**), and those with intensities higher than Apt2/Delta and lower than Scr/WT (green circles in **A**), were identified as potential WT and Delta specific binders, respectively. Their Omicron$_{XBB}$ binding signals were subsequently retrieved and summarized in a histogram (**B**), with red and green bins denoting WT- and Delta-selected candidates, respectively. This analysis suggested that WT- and Delta-specific candidates were present in the pool and remained largely unbound to Omicron$_{XBB}$, suggesting a good selectivity against Omicron$_{XBB}$.

**Supplementary Figure 6**

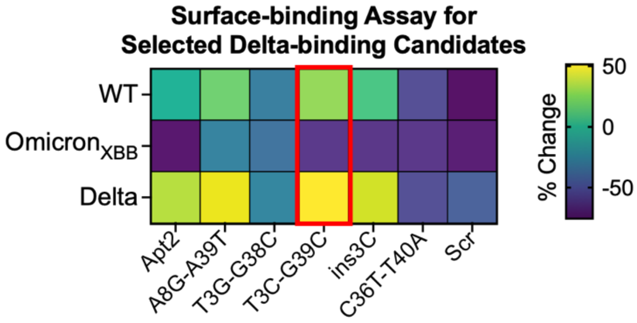

**Fig. S6. Use streptavidin-mediated surface binding assay to quickly examine the binding properties of the selected Delta-binding candidates.** Five candidates were selected from the on-chip results, as described in the main text. Together with the original Apt2 and Scr, the surface binding assay described in **fig. S1B** was performed to quickly examine the binding properties. The percentage changes were calculated based on the intensity of Apt2/WT. This experiment showed that most Delta-binding candidates have similar binding profiles to Apt2, with T3C-G39C double mutant (highlight in a red box) exhibiting the strongest binding to Delta. This agreed with the ELONA results showed in **Fig. 3C**.

**Supplementary Figure 7**

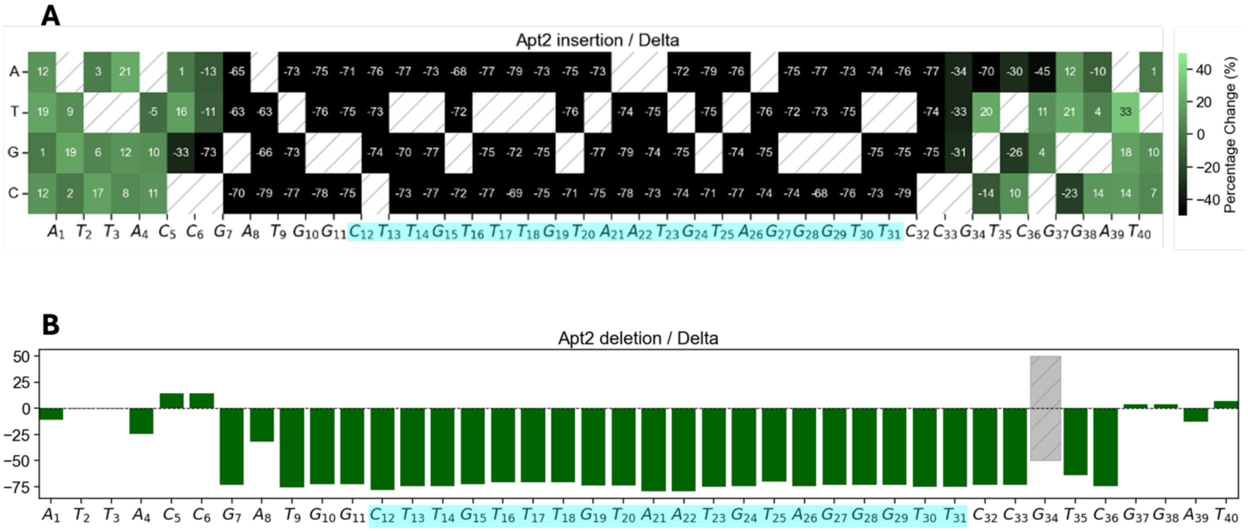

**Fig. S7. Single-insertion heatmaps and deletion bar plot for the *MiSeq* screening for SARS-CoV-2 Delta spike protein. (A)** Single-insertion heatmap. **(B)** Deletion bar plot. Together with the single-mutation results (**Fig. 3A**), these data showed that the loop motif remained significantly conserved for Delta binding. Any base substitution within the loop completely abolished binding. That is why we chose to focus on studying the variations in the stem (**Fig. 3B**). The missing data point (deletion at G34) is shown in grey. The loop motif is highlighted in cyan. Numbers represent the mean fluorescence intensity percentage change. Refer to **Methods** for more details.

**Supplementary Figure 8**

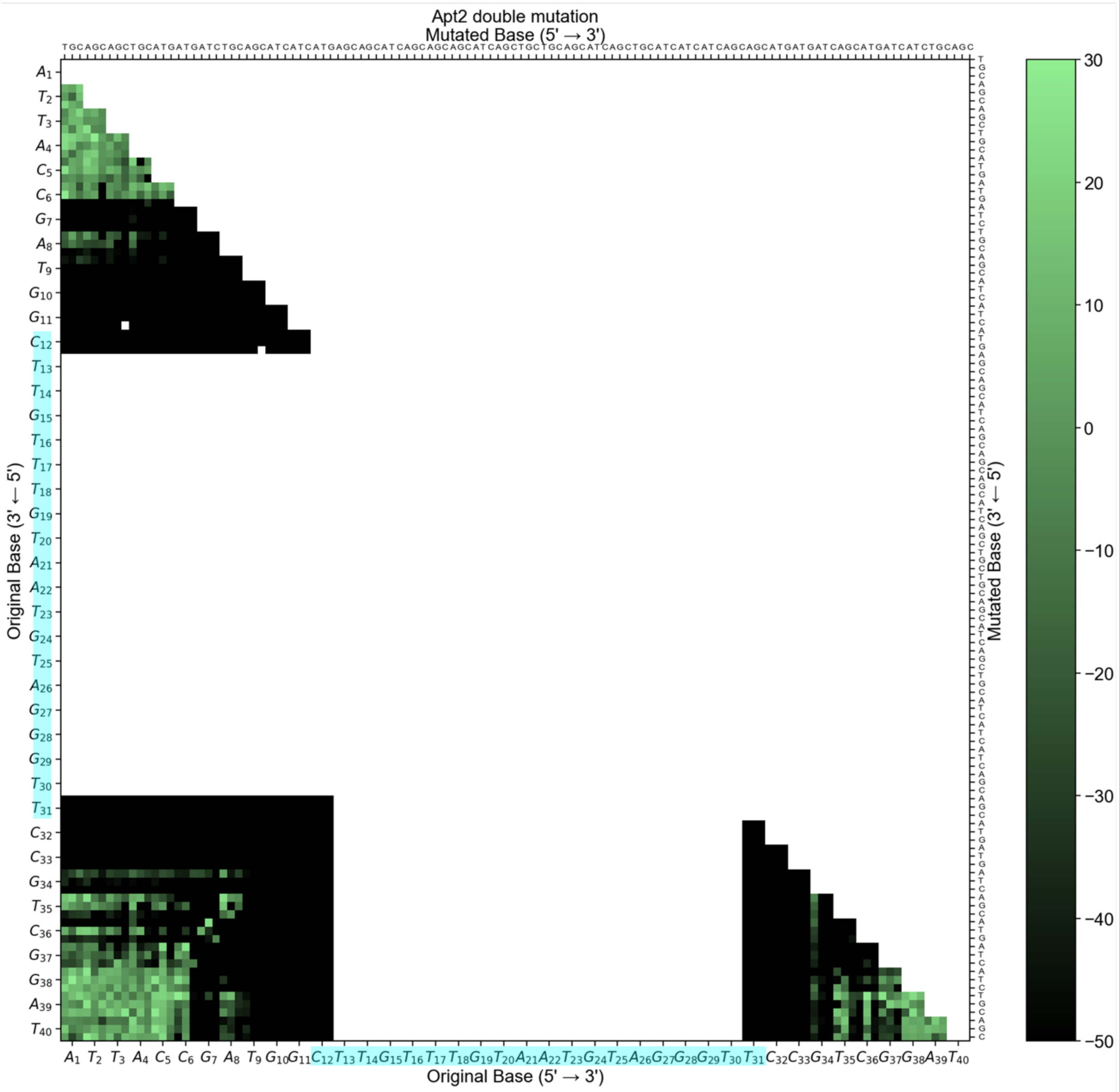

**Fig. S8. *MiSeq* screening-generated double-mutation heatmap (stem only) for Delta variant.** The data, together with **fig. S7**, showed that not only the loop motif, but bases close to the loop motif (i.e., G10 to C12; T31 to C33) also remained significantly conserved for Delta binding. Any base substitution there completely abolished binding. Missing data points were left blank. The loop motif is highlighted in cyan. Numbers represent the mean fluorescence intensity percentage change. Refer to **Methods** for more details.

**Supplementary Figure 9**

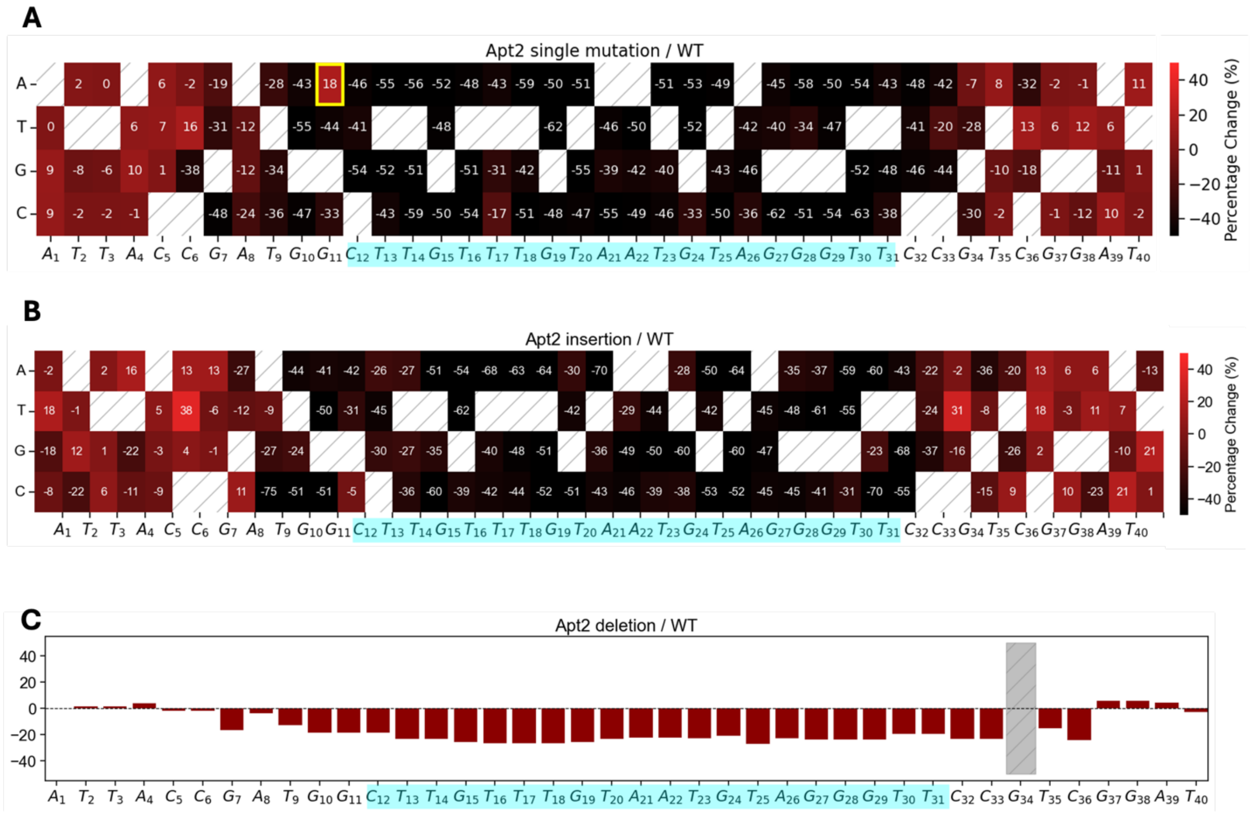

**Fig. S9. Single-mutation, single-insertion heatmaps and deletion bar plot for the *MiSeq* screening for SARS-CoV-2 WT spike protein. (A)** Single-mutation heatmap. **(B)** Single-insertion heatmap. **(C)** Deletion bar plot. These data showed that the loop motif was significantly conserved for WT binding, while G11A is an exception (highlight in a yellow box). The missing data point (deletion at G34) is shown in grey. The loop motif is highlighted in cyan. Numbers represent the mean fluorescence intensity percentage change. Refer to **Methods** for more details.

### Supplementary Figure 10

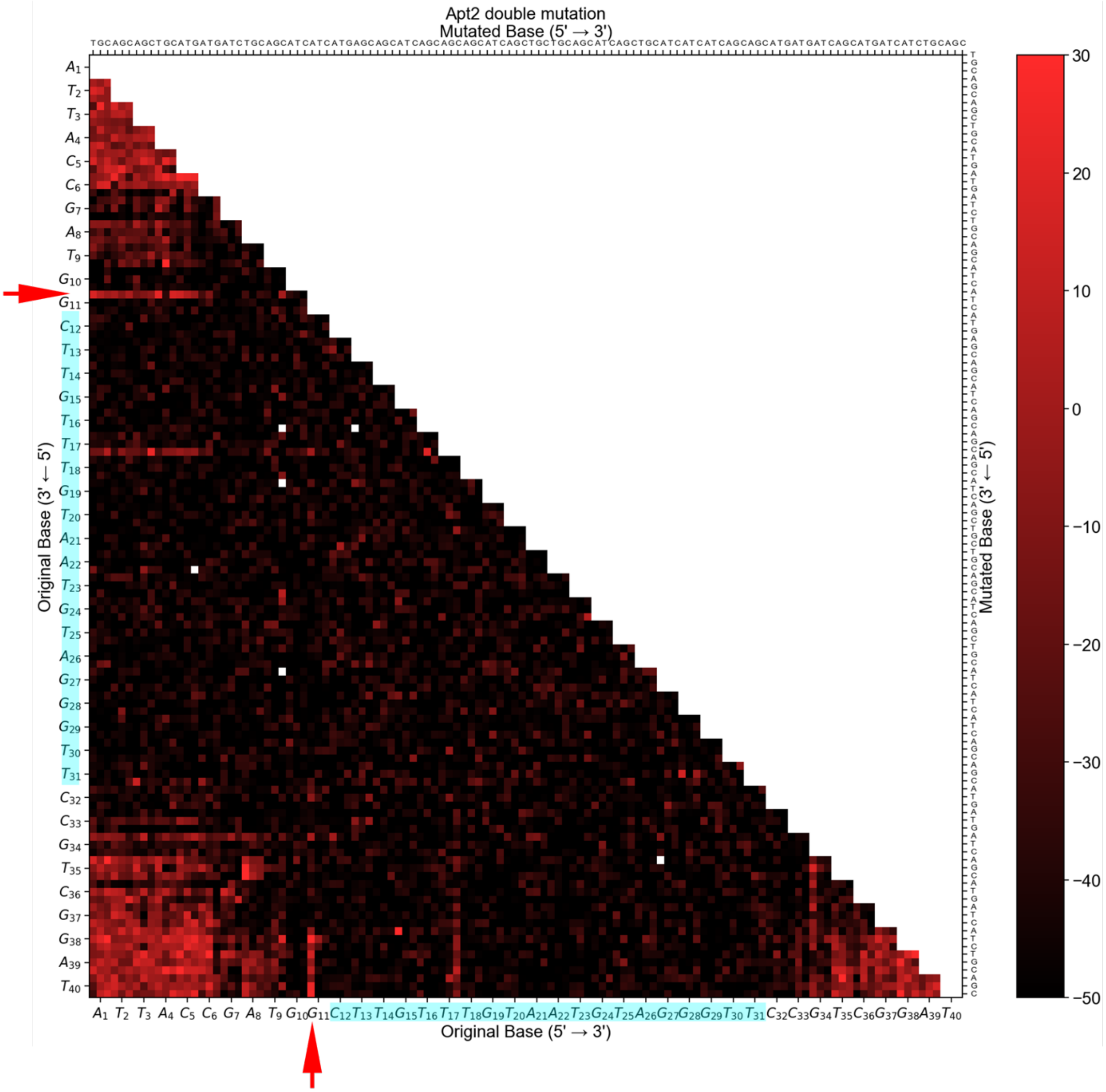

**Fig. S10. *MiSeq* screening-generated double-mutation heatmap for WT variant.** Double mutants containing G11A were indicated with two red arrows, indicating that this substitution is tolerated even adjacent to highly conserved bases. Missing data were left blank. The loop motif is highlighted in cyan. Numbers represent the mean fluorescence intensity percentage change. Refer to **Methods** for more details.

### Supplementary Figure 11

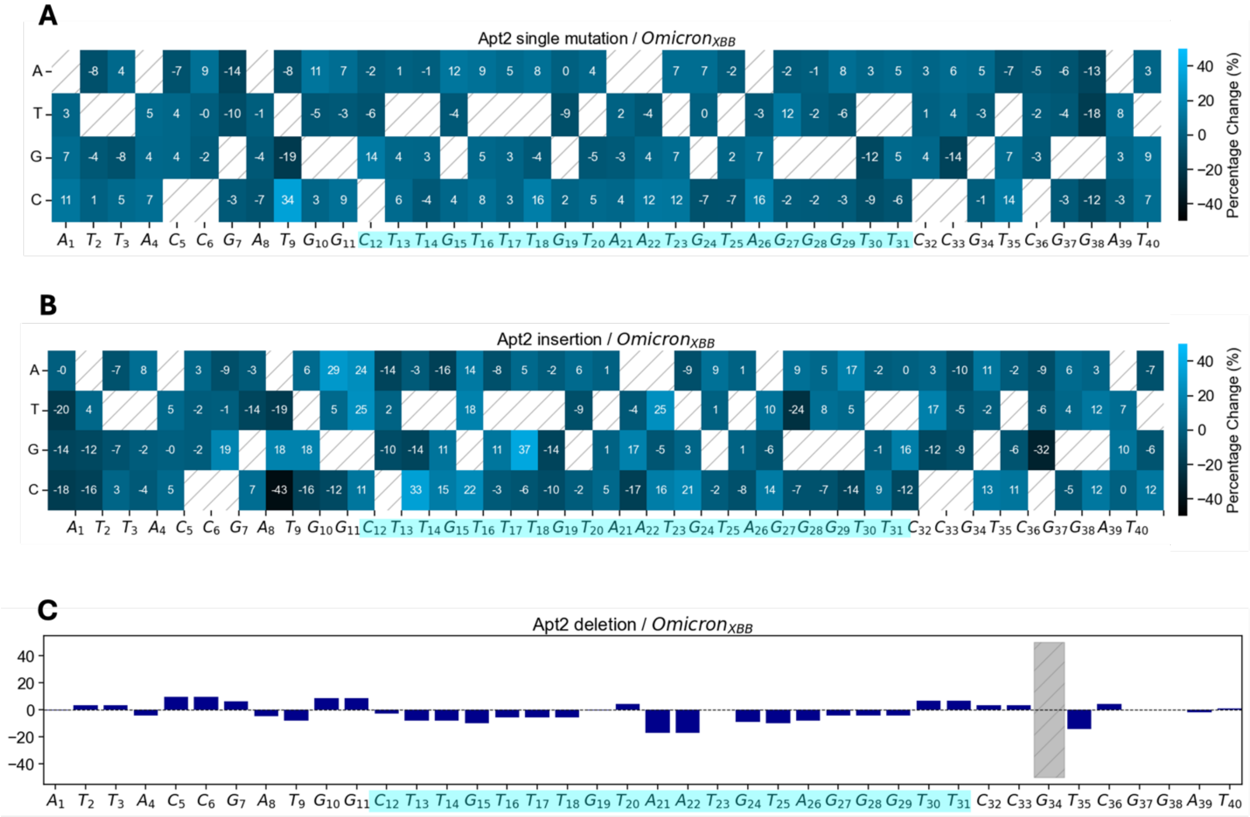

**Fig. S11. *MiSeq* screening-generated single-mutation, double-mutation heatmaps and deletion bar plot for Omicron$_{\text{XBB}}$ variant.** (**A**) Single-mutation heatmap. (**B**) Single-insertion heatmap. (**C**) Deletion bar plot**.** These analyses did not provide useful information for identifying Omicron$_{\text{XBB}}$ binders. Alternative approaches to mutating Apt2 are therefore required (**Fig. 4A-B**). The missing data point (deletion at G34) is shown in grey. The loop motif is highlighted in cyan. Numbers represent the mean fluorescence intensity percentage change. Refer to **Methods** for more details.

### Supplementary Figure 12

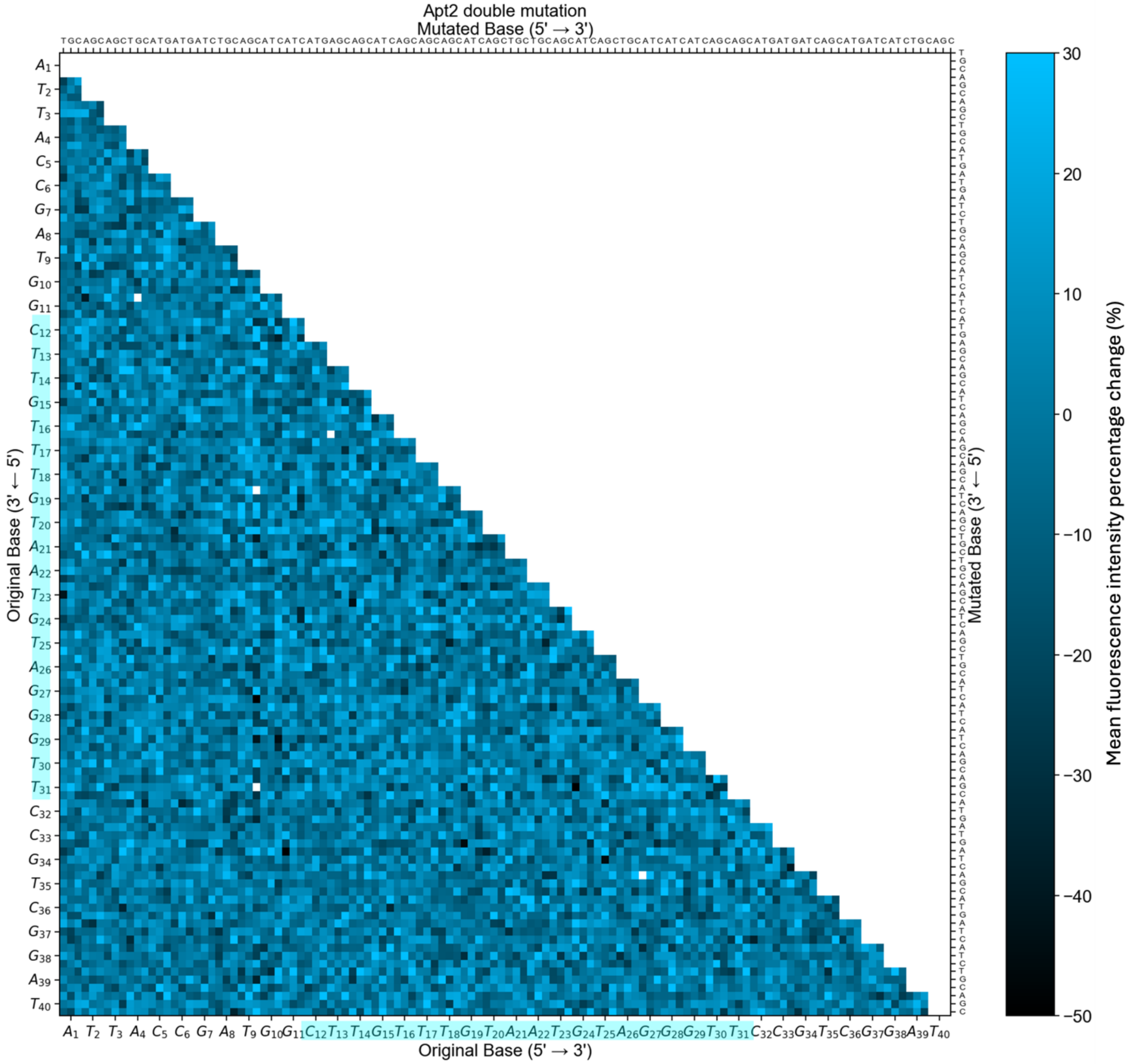

**Fig. S12. *MiSeq* screening double-mutation heatmap for Omicron$_{XBB}$ variant.** This analyses, together with the analyses in **fig. S11**, did not provide useful information for identifying Omicron$_{XBB}$ binders. Alternative approaches to mutating Apt2 are therefore required (**Fig. 4A-B**). The missing data points were left blank. The loop motif is highlighted in cyan. Numbers represent the mean fluorescence intensity percentage change. Refer to **Methods** for more details.

**Supplementary Figure 13**

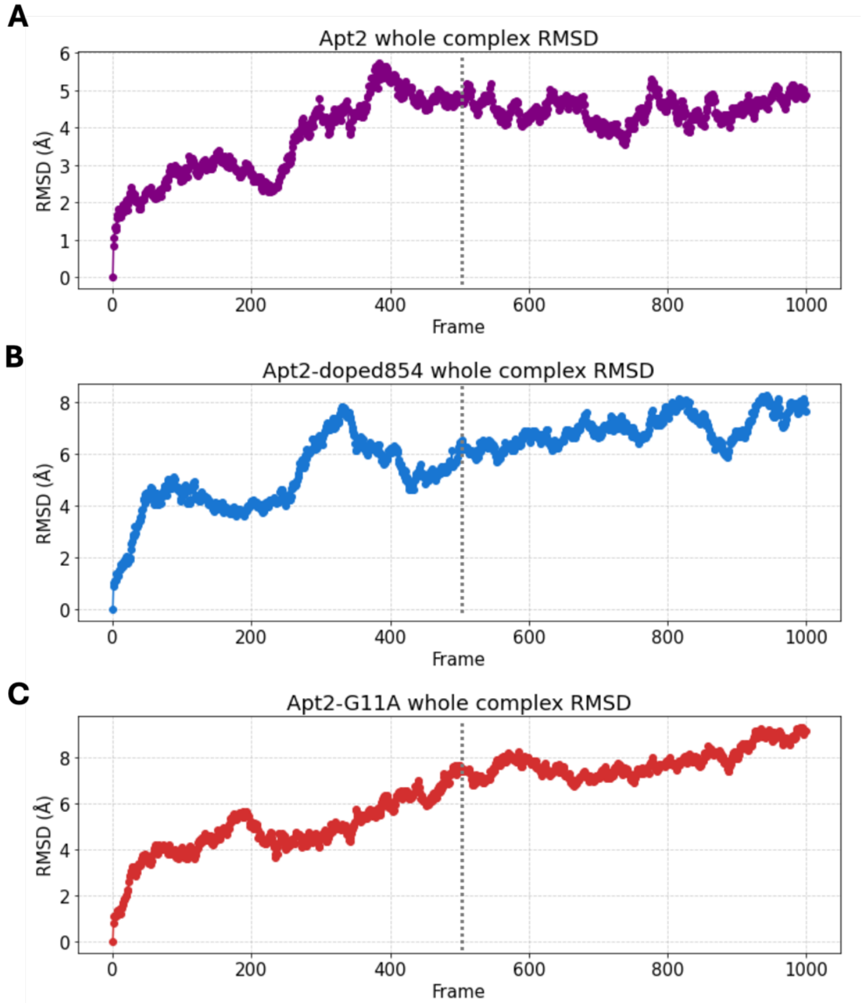

**Fig. S13. Root mean square deviation (RMSD) of the whole structures during MD simulation.** (**A**) Apt2. (**B**) doped854. (**C**) G11A. Frames were collected every 10 ps within the 10-ns simulation. RMSDs were computed with respect to the initial structure (frame 0). A rapid increase in RMSD was observed during the first ~5 ns (dotted lines), consistent with structural relaxation and non-equilibrated dynamics [16]. Between 5 and 10 ns, the RMSD exhibited a plateau, indicating that the system had reached equilibrium. Therefore, analyses were performed using only the second 5- ns period of the simulations.

### Supplementary Figure 14

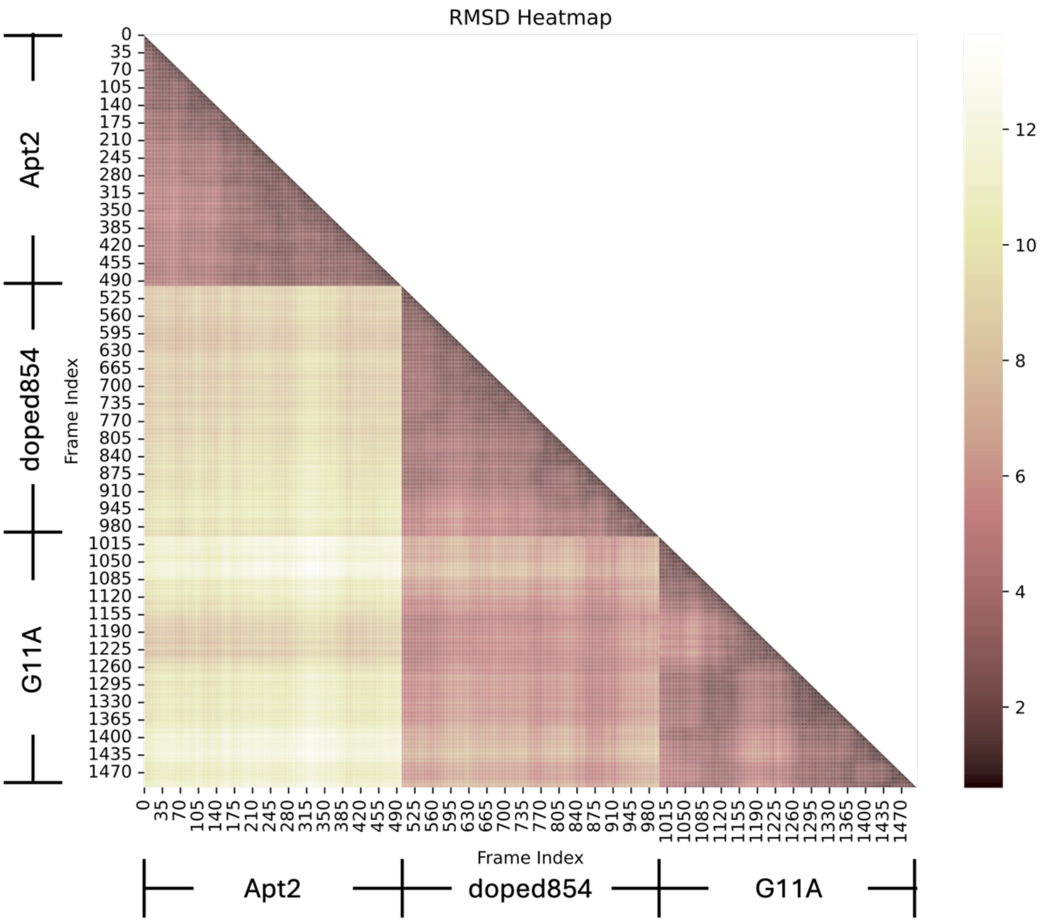

**Fig. S14. Cross validation within the second 5-ns period (right-hand side of the dash line) in fig. S13 MD simulations,** showing that group-group variance is much higher than in-group variance, suggesting acceptable dynamics simulations [17]. The in-group RMSD matrix was further used for clustering the dominant structures for each mutant (**Methods**).

**Supplementary Figure 15**

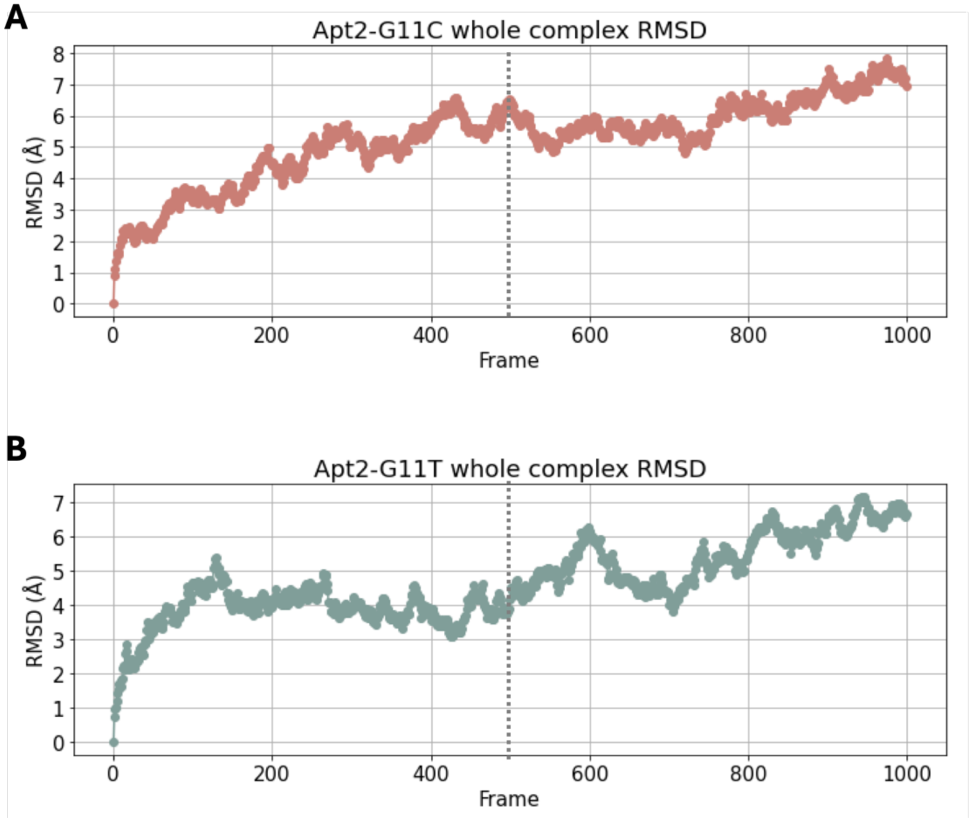

**Fig. S15. Root mean square deviation (RMSD) of the whole structures during MD simulation.** (**A**) G11C. (**B**) G11T. Frames were collected every 10 ps within the 10-ns simulation. RMSDs were computed with respect to the initial structure (frame 0). A rapid increase in RMSD was observed during the first ~5 ns (dotted lines), consistent with structural relaxation and non-equilibrated dynamics. Between 5 and 10 ns, the RMSD exhibited a plateau, indicating that the system had reached equilibrium. Therefore, analyses were performed using only the second 5-ns period of the simulations.

**Supplementary Figure 16**

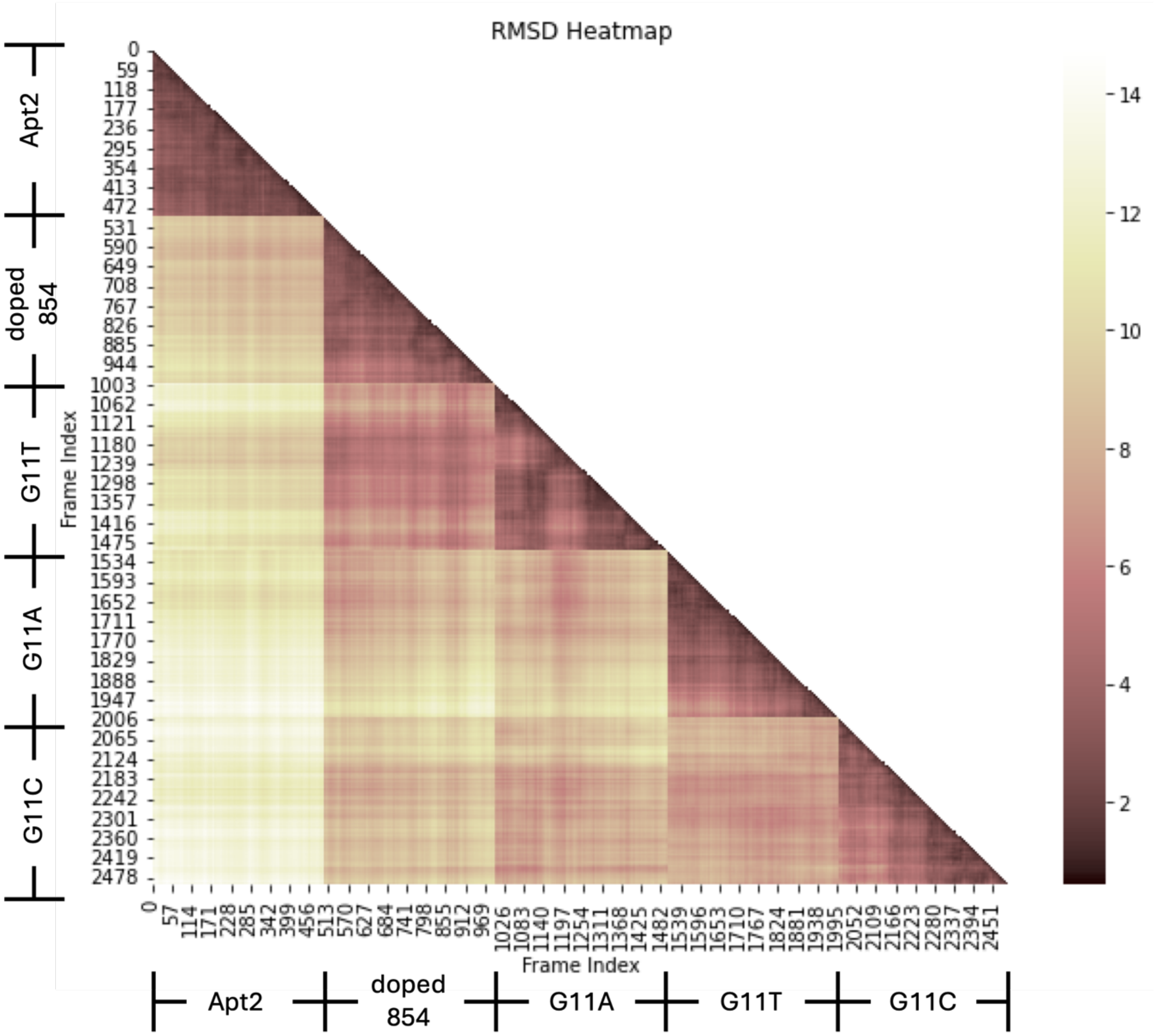

**Fig. S16. Cross validation within latter 5 ns,** showing that group-group variance is much higher than in-group variance, including G11C and G11T. The data suggested acceptable dynamics simulations.

### Supplementary Figure 17

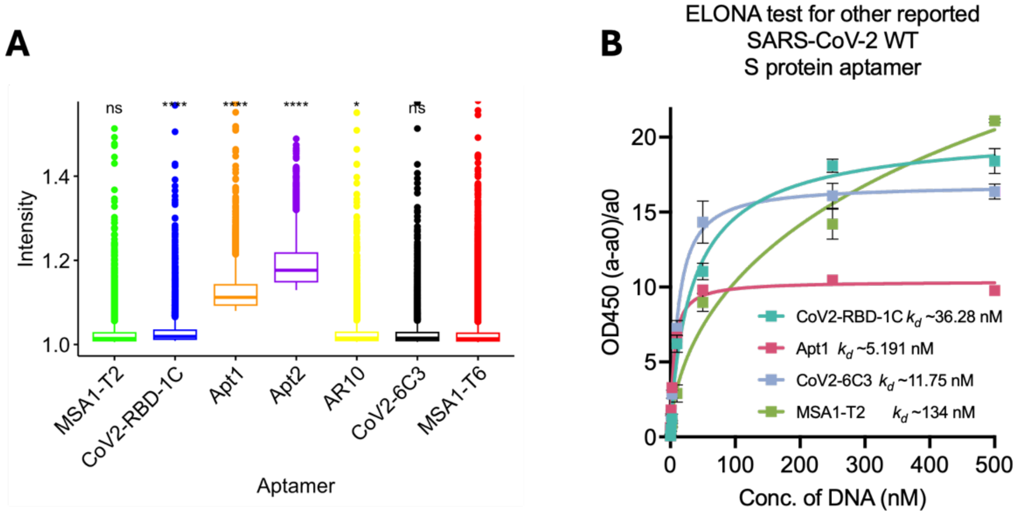

**Fig. S17. Initial trials on other published aptamers. (A)** *MiSeq* fluorescence intensity of various published SARS-CoV-2 spike protein aptamers when adding WT. MSA1-T6 is a non-binder to SARS-CoV-2 spike protein according to Li group's report.[2] Sample sizes: MSA1-T2: 26,344; CoV2-RBD-1C: 29,372; Apt1: 33,986; Apt2: 25,611; AR10: 33,986; CoV2-6C3: 29,234; MSA1-T6: 99,655. **(B)** Characterize some of these aptamers with ELONA. ANOVA test compared to the negative control sequence (MSA1-T6) on the chip: *, $p < 0.05$; **, $p < 0.01$; ***, $p < 0.001$; ****, $p < 0.0001$; ns, $p > 0.05$. Sources and the sequences of these aptamers can be found in **Supplementary Table S1**. This is why we chose Apt2 as our model system.

**Supplementary Figure 18**

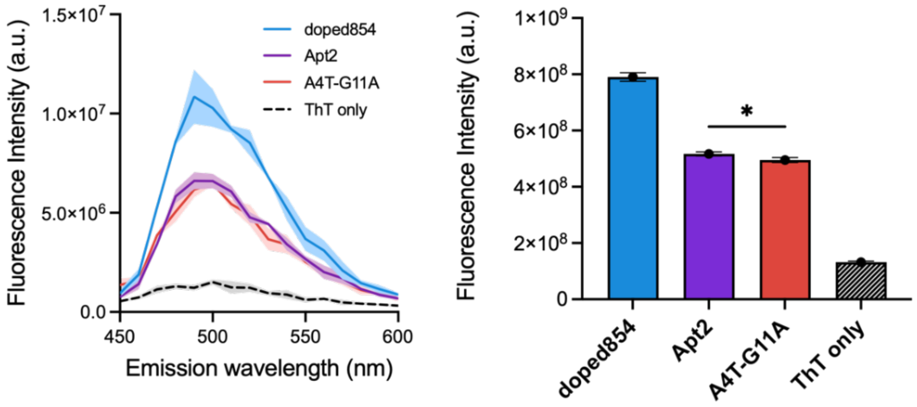

**Fig. S18. Fluorescence activation of thioflavin T by DNA strands (i.e., ThT assay),** comparison among doped854, Apt2 and A4T-G11A. Two-tailed unpaired t test was performed to compare the fluorescence intensities between A4T-G11A and Apt2, leading to a P-value of 0.0297. ThT assay is commonly used to detect the G quadruplex structure in nucleic acids [18]. Each sample has 3 duplicates, and the shadings indicate standard deviation. Fluorescence emission spectra were measured using a plate reader (SpectraMax i3, Molecular Devices) with step size of 5 nm.

**Supplementary Figure 19**

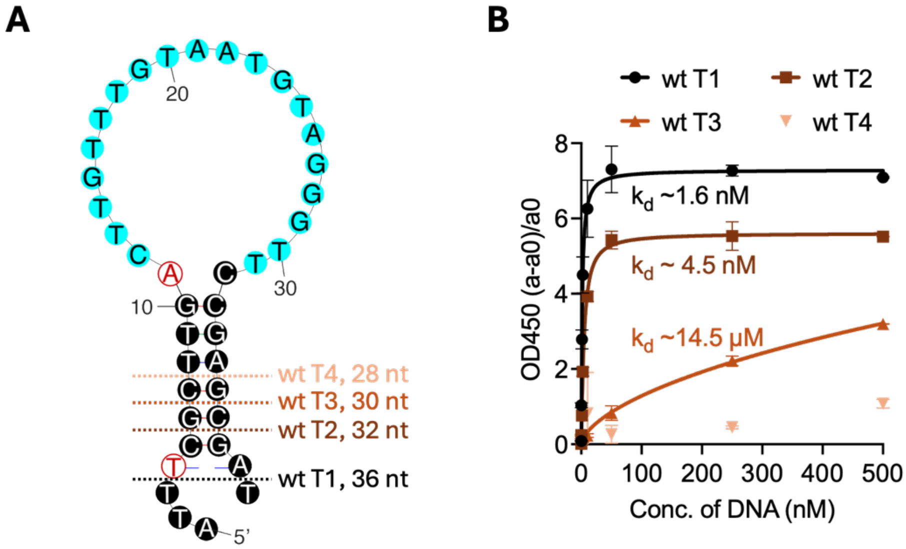

**Fig S19. Truncation trials with ELONA.** (**A**) Different truncation sites on A4T-G11A with a scrambled lower stem. (**B**) ELONA results for all truncated aptamers showed in (**A**). According to the results, wt T1 retained the strongest binding to WT. Therefore, wt sensor was built on it (**Fig. 7A** and **Supplementary Table S1**).

**Supplementary Figure 20**

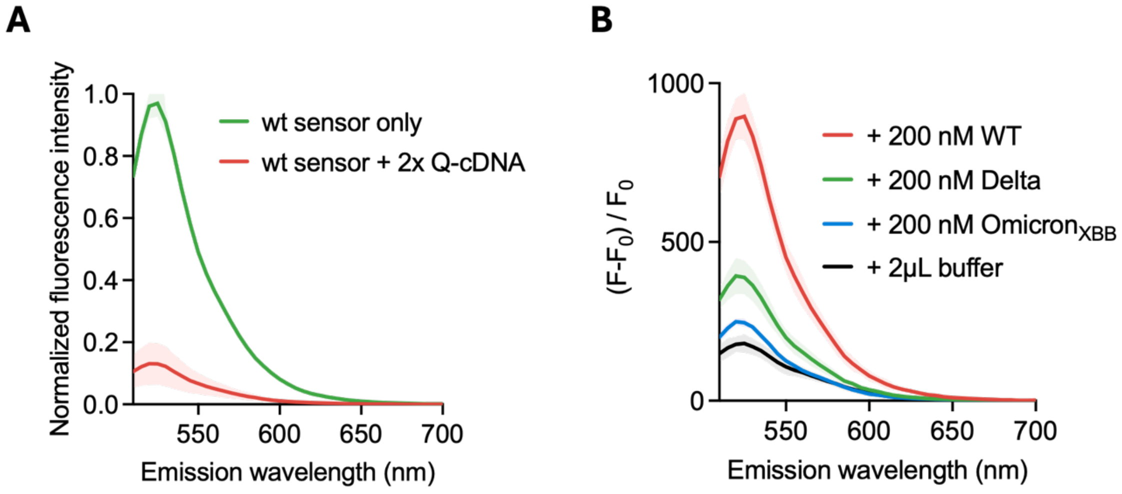

**Fig S20. Quenching efficiency test and single-point comparison for different spike protein strains for the strand-displacement fluorescent sensor.** (**A**) After adding 100 nM Q-cDNA to 50 nM wt sensor, the fluorescence was quenched by 86%. Sensor design can be found in **Fig. 7A**. (**B**) Upon addition of 200 nM WT spike protein to the sample, fluorescence was enhanced by ~3 fold in AUC (area under curve), significantly exceeding the responses observed for Delta (~1.9 fold) and Omicron$_{XBB}$ (~1.2 fold) at the same concentration. For both (**A**) and (**B**), the lines represent the mean values, while the shadings represent the standard deviations of two duplicates. Refer to **Methods** for more details.

**Supplementary Figure 21**

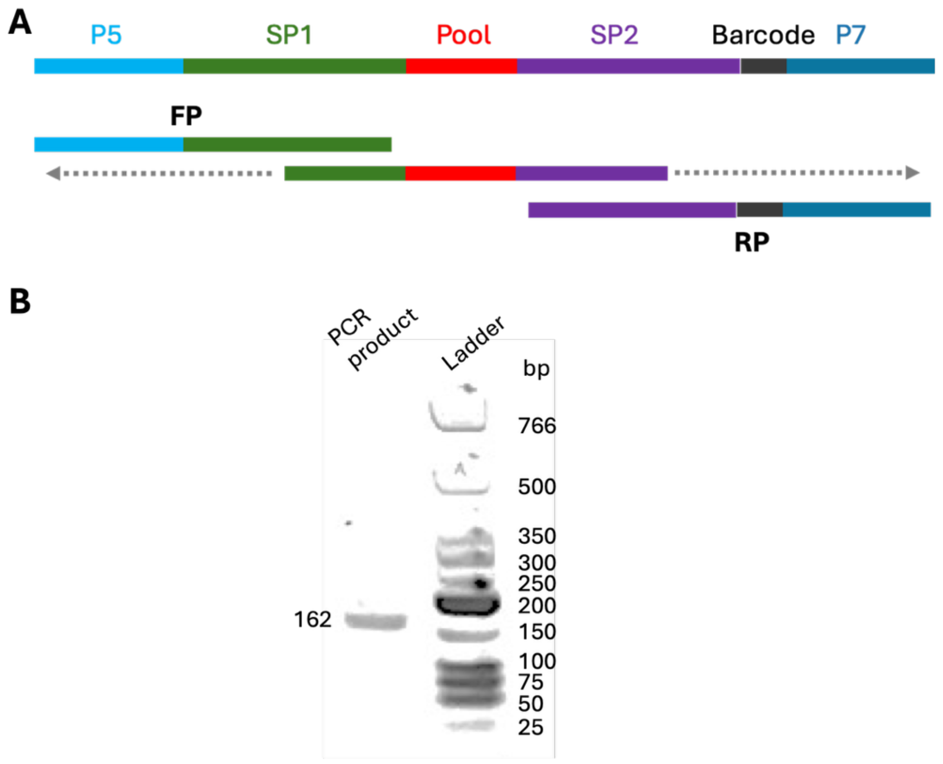

**Fig. S21. Construct the sequences for NGS flow using overlap extension PCR.** (**A**) Illustration of the final PCR product and the process. Sequences can be found in **Supplementary Table S1**. (**B**) 2% agarose gel showed a clear single band at a desired position, suggesting a successful and clean extension. The gel was imaged with Cytiva Typhoon RGB Imager.

**Movies S1-S3 (Separate files)**

**Apt2, G11A, doped 854's molecular dynamics in latter 5 ns during simulations**

One frame was extracted every 200 frames to reduce the size of the trajectory file. Color code: tails-green; lower stem: yellow; upper stem: magenta; loop (motif): cyan. These movies provide a qualitative visualization of base-level dynamics throughout the latter 5-ns simulations, complementing the quantitative RMSF analysis shown in **Fig. 6**.

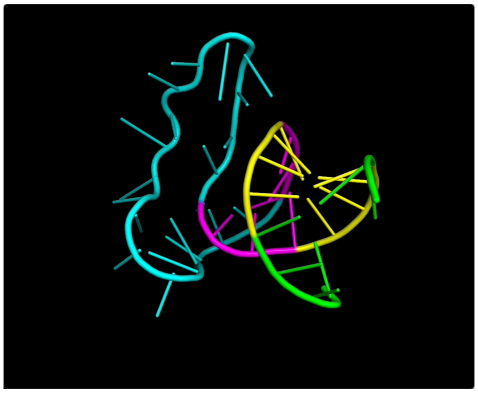

Screenshot for Movie S1

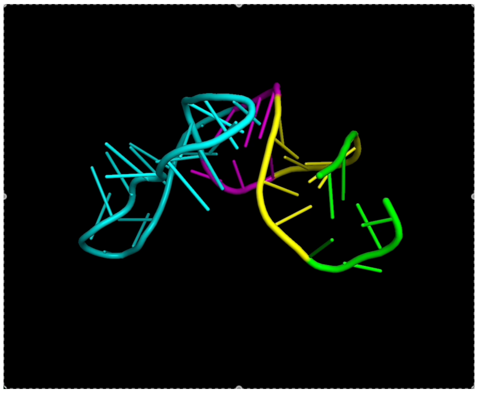

Screenshot for Movie S2

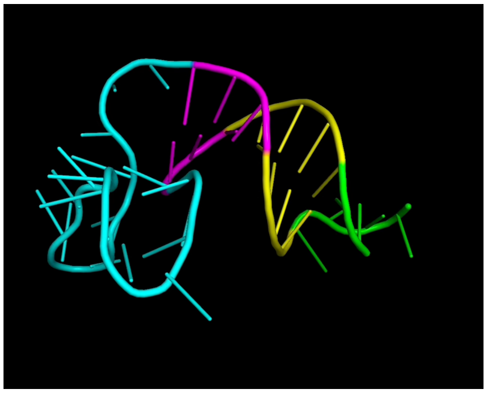

Screenshot for Movie S3
